# Intermediate basal cell population in prostate homeostasis and cancer initiation

**DOI:** 10.1101/2023.05.12.540502

**Authors:** Wangxin Guo, Xiaoyu Zhang, Lin Li, Pengfei Shao, Hongjiong Zhang, Kuo Liu, Chao Liang, Shuoming Wang, Yunyi Peng, Yi Ju, Chen Yu, Luonan Chen, Bin Zhou, Dong Gao

**Affiliations:** State Key Laboratory of Cell Biology, Shanghai Key Laboratory of Molecular Andrology, Shanghai Institute of Biochemistry and Cell Biology, Center for Excellence in Molecular Cell Science, Chinese Academy of Sciences, Shanghai 200031, China; University of Chinese Academy of Sciences, Beijing 100049, China; Department of Urology, First Affiliated Hospital of Nanjing Medical University, Nanjing, China; Institute of Cancer Research, Shenzhen Bay Laboratory, Shenzhen 518132, China; School of Life Science and Technology, ShanghaiTech University, Shanghai 201210, China; Key Laboratory of Systems Health Science of Zhejiang Province, School of Life Science, Hangzhou Institute for Advanced Study, University of Chinese Academy of Sciences, Hangzhou 310024, China

## Abstract

Many glandular epithelia are mainly composed of basal cells and luminal cells, including the prostate gland. Adult prostate basal and luminal cells are independently self-sustained by unipotent stem cells that can reactivate multipotency under prostate inflammation and carcinogenesis contexts. However, the defined basal stem cell populations responsible for prostate regeneration and their cell fates in prostate homeostasis, inflammation and carcinogenesis remain unclear. Using a genetic proliferation tracer (ProTracer) system, we found that basal cells exhibited extensive cell loss and proliferation during androgen-mediated prostate regression and regeneration, respectively. A rare intermediate basal cell population that expresses luminal cell markers (*Nkx3*.*1* and *Pbsn*) (termed Basal-B) and a large basal cell population (termed Basal-A) were identified in mouse prostates by single-cell RNA sequencing. Basal-B cells exhibited a greater capacity for organoid formation and luminal cell differentiation *in vitro*. Genetic lineage tracing using dual recombinases showed that prostate homeostasis and regeneration are not driven by specific basal cell types. Fate-mapping results showed that Basal-B cells had a greater tendency to generate luminal cells under bacteria-induced prostate inflammation. Deletion of *Pten* in basal cells resulted in Basal-A-to-Basal-B-to-luminal transition and prostatic intraepithelial neoplasia. Moreover, the human Basal-B-cell population was significantly increased in human benign prostate hyperplasia and prostatic intraepithelial neoplasia samples compared with normal prostate samples. This study identifies intermediate Basal-B cells as a potential stem cell population and provides genetic evidence of prostate basal cell lineage plasticity under physiological and pathological contexts.

## Introduction

Organ homeostasis and repair are often driven by tissue-specific stem cell populations (*1, 2*). Lineage-restricted stem cells can have remarkable lineage plasticity and long-term self-renewing capacities under physiological and pathological conditions (*3*). The normal adult prostate epithelium is mainly composed of basal cells and luminal cells (*4, 5*). Isolated basal cells can generate both basal cells and luminal cells under *in vivo* prostate cell transplantation (*6-9*) and *in vitro* organoid formation assays (*10-12*). Genetic lineage tracing results demonstrated that adult prostate basal cells contain lineage-restricted unipotent progenitor cells during prostate homeostasis and androgen-induced prostate regeneration (*13-15*). Adult unipotent prostate basal stem cells can reactivate multipotency under prostate inflammation, luminal cell ablation and carcinogenesis contexts (*14, 16-18*). However, distinct prostate basal stem cells with defined markers are still unknown. It is clear that the majority of luminal cells are androgen dependent and undergo apoptosis following androgen deprivation therapy by surgical or pharmacological castration (*19-21*). However, the basal cell fates under androgen deprivation (Fig. 1A), prostate inflammation and carcinogenesis conditions remain undefined because fate mapping of basal cells has only been performed on a few unclassified basal cells.

**Fig. 1.**
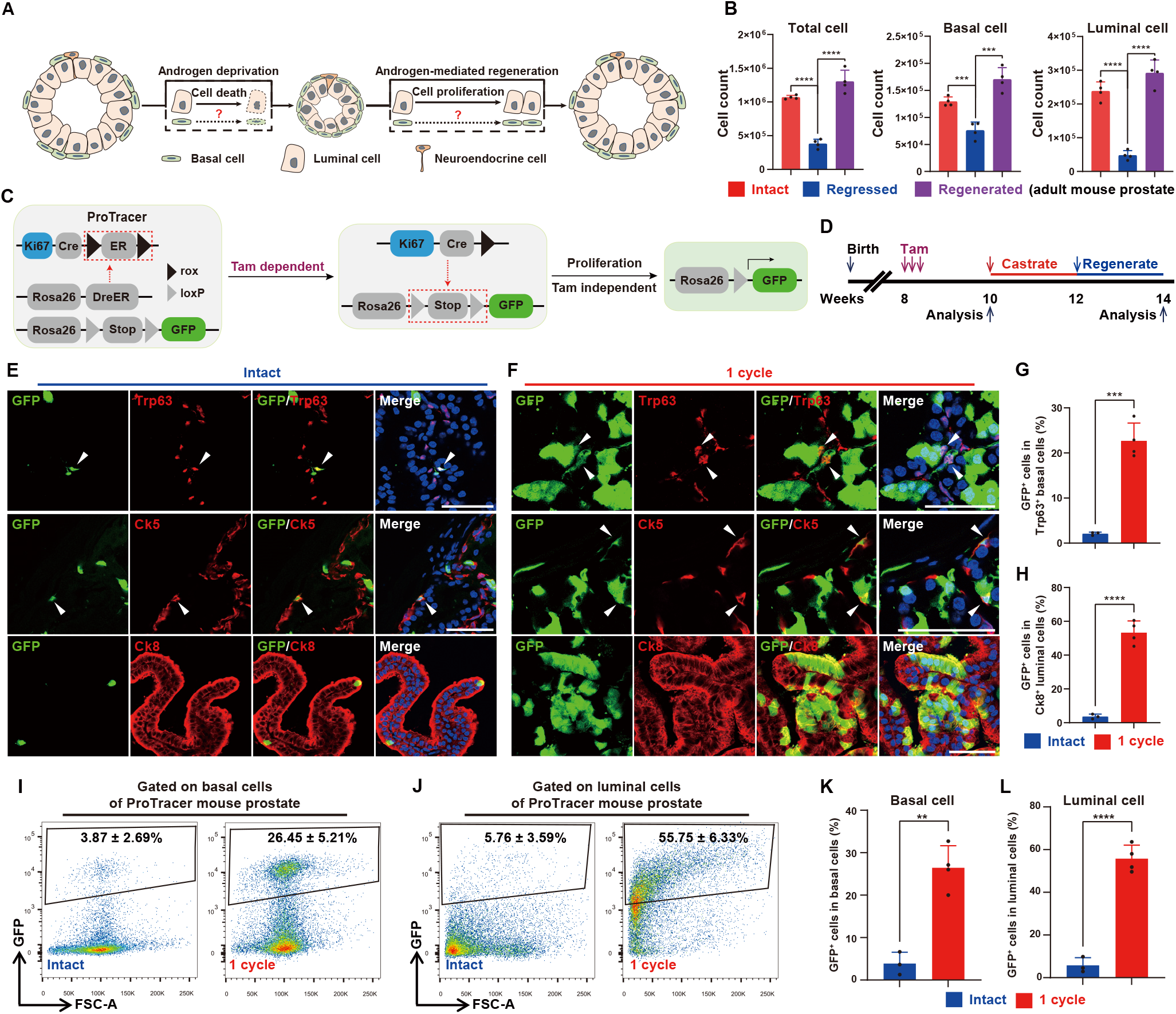
Extensive basal cell loss and proliferation during androgen-mediated prostate regression and regeneration. (**A**) Schematic diagram illustrating that basal cell fates remain undefined during prostate regression and regeneration. (**B**) Bar graphs showing the cell number of total, basal and luminal cells in intact, regressed and regenerated adult mouse prostates (*n* = 4 mice, per group). (**C**) Construction strategy of ProTracer (*R26-DreER;Ki67-CrexER;R26-LSL-GFP*) mice for cell proliferation tracing. (**D**) Schedule of tracing proliferated cells during prostate regression-regeneration in ProTracer mice. Tam is an abbreviation for tamoxifen. (**E** and **F**) Immunofluorescence staining of GFP, Trp63, Ck5 and Ck8 in intact (E) and regenerated mouse prostates after 1 cycle of prostate regression-regeneration (F) in ProTracer mice. White arrows indicate GFP^+^ proliferated basal cells. (**G** and **H**) Bar graphs showing the percentage of GFP^+^ cells in Trp63^+^ basal cells (G) or Ck8^+^ luminal cells (H) of intact (*n* = 3 mice) and regenerated (*n* = 4 mice) mouse prostates in ProTracer mice after 1 cycle of prostate regression-regeneration. (**I** and **J**) FACS analysis showing the percentage of GFP^+^ cells in basal cells (I) or luminal cells (J) of intact (*n* = 3 mice) and regenerated (*n* = 4 mice) mouse prostates in ProTracer mice after 1 cycle of prostate regression-regeneration. (**K** and **L**) Bar graphs showing the percentage of GFP^+^ cells in basal (K) or luminal (L) cells of intact (*n* = 3 mice) and regenerated (*n* = 4 mice) mouse prostates in ProTracer mice after 1 cycle of prostate regression-regeneration. Scale bars, 50 m. Data are shown as the mean ± SD. Data were analyzed by unpaired two-tailed Student’s t test; \*\**p* < 0.01, ^***^*p* < 0.001 and ^****^*p* < 0.0001.

An intermediate cell population (coexpressing basal and luminal cell markers) enriched in areas of human prostate inflammatory atrophy (PIA) is a suspected target for prostate carcinogenesis (*17, 22*). Uropathogenic *E. coli* infection induced the expansion of the intermediate cell population and basal-to-luminal cell transition in mouse prostates (*16*). Lineage tracing using *Ck14-CreER* demonstrated that the intermediate cells apparently arose from CK14-expressing basal cells. Exemplified mixed basal-luminal cells have been identified in advanced prostate cancer (*23-26*). Single-cell RNA sequencing (scRNA-seq) technology further highlighted the unexpected cell lineage complexity in normal prostate and prostate cancer tissues (*27-32*). However, the defined cell identity of prostate intermediate cell populations and their lineage plasticity in prostate diseases are still unknown.

Investigating cell lineage plasticity provides fundamental information about the role of stem cells in tissue development, regeneration and diseases. Genetic lineage tracing techniques, such as the Cre-*loxP* system, are powerful tools to study the cell lineage plasticity of stem cells or progenitors (*2*). However, the precision of these genetic systems heavily relies on the recombinase expression specificity in targeted stem cells. Furthermore, these lineage tracing systems might introduce a potential bias, which only focuses on marker-labeled cells rather than the entire cell population. Due to these limitations, genetic lineage tracing studies have led to conflicting conclusions concerning basal and luminal cell fates in prostate regeneration and the cell origin of prostate cancer (*13-15, 21, 33-35*).

Here, we simultaneously characterized prostate basal, luminal and intermediate cell populations during prostate homeostasis, regeneration, inflammation and cancer initiation using a genetic proliferation tracer (ProTracer) system (*36*), enhanced lineage tracing strategies that use dual recombinases (Cre and Dre) (*37*) and scRNA-seq technology (*38*).

## Results

### Extensive basal cell loss and proliferation during prostate regression and regeneration

Androgen deprivation therapy is the cornerstone for prostate cancer treatment. Previous studies have shown that androgen deprivation is often linked to the loss of luminal cells with high androgen receptor (AR) expression levels, but basal cells (AR^low^) are largely unaffected (*19-21*). However, basal cell fates under androgen deprivation need further investigation (Fig. 1A) because in addition to AR expression levels, many other factors may also affect the cell fates of basal cells (*14*). To ascertain the response of basal cells to androgen deprivation *in vivo*, we performed prostate regression-regeneration assays using castration and testosterone administration in C57BL/6 (B6) male mice and compared the cell number of basal cells (Lin^-^/Epcam^high^/CD49f^high^) and luminal cells (Lin^-^/Epcam^high^/CD49f^low^) within the prostates by fluorescence-activated cell sorting (FACS) (fig. S1, A to C). As expected, there was dramatic luminal cell loss (79.2 ± 8.45%) after castration and luminal cell expansion during prostate regeneration (Fig. 1B and fig. S1D). Surprisingly, basal cells also had extensive cell loss (40.73 ± 14.73%) after castration and cell expansion during prostate regeneration (Fig. 1B and fig. S1D).

To directly record cell type-specific proliferation during prostate regeneration *in vivo*, we used the genetic proliferation tracer system ProTracer (*36*) (*R26-DreER*;*Ki67-Cre-rox-ER-rox*;*R26-loxp-stop-loxp-GFP*) to seamlessly record the proliferation events of entire prostate cell populations (Fig. 1C). Briefly, tamoxifen (Tam) treatment-induced DreER-rox recombination converts the coding sequence of inducible *Ki67-CrexER* into constitutively active *Ki67-Cre* DNA in DreER-expressing cells and permanently records active Ki67 transcription by irreversibly activating the *R26-GFP* reporter (Fig. 1C) (*36*). We euthanized mice, collected prostate samples and examined in situ prostate basal and luminal cell proliferation before castration and after one cycle of prostate regression-regeneration (Fig. 1D). Immunostaining for GFP, Trp63, Ck5 and Ck8 showed that both basal cells and luminal cells exhibited extensive cell proliferation during prostate regeneration (Fig. 1, E and F). Quantification of the percentage of GFP^+^ cells before and after one cycle of prostate regression-regeneration revealed that the approximate proliferation rates of basal cells and luminal cells were 0.206 (from 0.021 ± 0.004 to 0.227 ± 0.039) and 0.497 (from 0.036 ± 0.015 to 0.533 ± 0.069) per cycle of prostat e regression-regeneration, respectively (Fig. 1, G and H). FACS analysis also showed that the proliferation of GFP^+^ basal cells (Lin^-^/Epcam^high^/CD49f^high^) and luminal cells (Lin^-^/Epcam^high^/CD49f^low^) was significantly increased after one cycle of prostate regression-regeneration (Fig. 1, I to L). The continuous proliferation recording of all adult prostate cell populations suggests that both basal cells and luminal cells have extensive cell proliferation during prostate regeneration.

To simultaneously elucidate the cell fates of both basal cells (Trp63^+^) and luminal cells (Ck8^+^) during androgen-mediated prostate regression-regeneration, we utilized enhanced lineage tracing strategies that use dual recombinases (Cre and Dre) (*37*). We combined *Trp63-DreER* and *Ck8-CreER* with the Rosa26 traffic light reporter *R26-TLR* (short for *R26-CAG-rox-stop-rox-ZsGreen-Insulater-CAG-loxp-stop-loxp-tdTomato*) (*39*) for lineage tracing of Trp63^+^ basal cells (ZsGreen^+^), Ck8^+^ luminal cells (tdTomato^+^) and Trp63^+^Ck8^+^ intermediate cells (ZsGreen^+^tdTomato^+^) (fig. S2, A and B). Immunofluorescence analysis showed that ZsGreen^+^ cells were strictly positive for the basal cell marker Trp63, and the labeling efficiency was 93.61 ± 1.93%. The tdTomato ^+^ cells were strictly positive for the luminal cell marker Ck8, and the labeling efficiency was 8.21 ± 1.90% (fig. S2, C to E). We did not detect any ZsGreen^+^tdTomato^+^ intermediate cells in adult mouse prostates (fig. S2, C and F). After one cycle of androgen-mediated prostate regression-regeneration, fate mapping showed that ZsGreen^+^ and tdTomato^+^ cells still strictly maintained their original identity as basal cells and luminal cells, respectively (fig. S2, G to I). These results reveal that adult basal and luminal cells are strictly self-sustained during androgen-mediated prostate regression and regeneration. Taken together, cell proliferation tracing and enhanced lineage tracing results suggest that adult prostate basal cells also have extensive cell loss and cell proliferation during androgen-mediated prostate regression and regeneration.

### Basal-B cells are potential adult prostate basal stem cells

To elucidate the prostate basal cell hierarchy, we extracted all basal cells from our mouse prostate scRNA-seq dataset (*27*) and performed unsupervised clustering using Seurat 3.0. A large basal cell population (Basal-A) and a rare basal cell population (Basal-B) were identified in mouse prostates (Fig. 2A and fig. S3). Both Basal-A cells and Basal-B cells highly expressed basal cell markers, such as *Trp63, Ck5* and *Ck14* (fig. S3A). We identified marker genes to classify Basal-A cells (*Ck17, Dlk2* and *Prelp*) and Basal-B cells (*Nkx3*.*1, Pbsn, C1rb, Fgl1* and *Mmp7*) (Fig. 2, B and C and fig. S3). Interestingly, *Nkx3*.*1, Pbsn, Fgl1* and *Mmp7* were defined as Luminal-B cell markers (*27*). To confirm the existence of this rare intermediate Basal-B cell population, we performed coimmunofluorescence for basal cell markers, such as Trp63 and Ck5, and the Basal-B marker Nkx3.1. We confirmed that Basal-B cells (Trp63^+^/Ck5^+^/Nkx3.1^+^) had a scattered distribution pattern in mouse prostates (fig. S4).

**Fig. 2.**
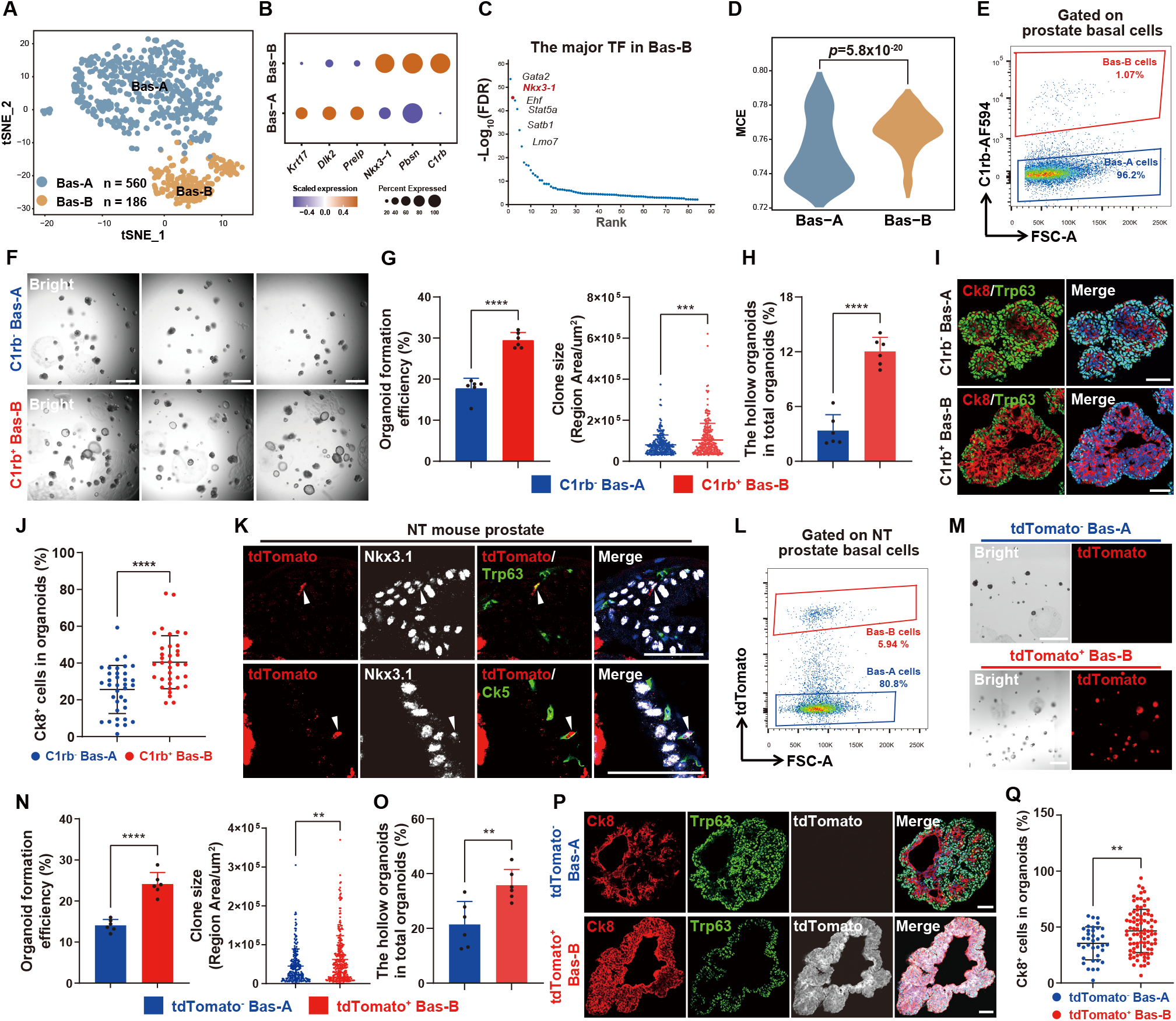
Basal-B cells are potential multipotent stem cells *in vitro*. **(A)** t-SNE plot shows clusters of Basal-A and Basal-B cells from adult mouse prostates. **(B)** Dot plot showing the expression levels of representative signature genes in Basal-A and Basal-B cells. **(C)** Visualization of significant differentially expressed transcription factors (TFs) in Basal-B cells. (**D**) Violin plot showing the MCE values of Basal-A and Basal-B cells. The *P* value is from a one-sided t test. (**E**) FACS plot showing C1rb^-^ Basal-A and C1rb^+^ Basal-B cells isolated from the basal cells (Lin^-^/Epcam^high^/CD49f^high^) of WT adult mouse anterior prostates. (**F**) Micrographs showing C1rb^-^ Basal-A and C1rb^+^ Basal-B-cell-derived organoids. Scale bar, 1 mm. (**G** and **H**) Bar graphs showing organoid formation efficiency, clone size (G) and percentage of hollow organoids (H) of C1rb^-^ Basal-A and C1rb^+^ Basal-B-cell-derived organoids. (**I**) Immunofluorescence staining of Ck8 and Trp63 in C1rb^-^ Basal-A and C1rb^+^ Basal-B-cell-derived organoids. Scale bars, 50 m. (**J**) Plot diagram shows the percentage of Ck8^+^ cells in C1rb^-^ Basal-A and C1rb^+^ Basal-B cell-derived organoids. (**K**) Immunofluorescence staining of Nkx3.1, Trp63, Ck5 and tdTomato in NT mouse prostates. White arrows indicate tdTomato^+^Nkx3.1^+^ Basal-B cells. Scale bars, 50 m. (**L**) FACS plot showing tdTomato^-^ Basal-A and tdTomato^+^ Basal-B cells isolated from the basal cells (Lin^-^/Epcam^high^/CD49f^high^) of NT mouse anterior prostates. (**M**) Micrographs showing tdTomato^-^ Basal-A and tdTomato^+^ Basal-B-cell-derived organoids. (**N** and **O**) Bar graphs show organoid formation efficiency, clone size (N) and percentage of hollow organoids (O) of tdTomato^-^ Basal-A and tdTomato^+^ Basal-B-cell-derived organoids. Scale bar, 1 mm. (**P**) Immunofluorescence staining of Ck8, Trp63 and tdTomato in tdTomato^-^ Basal-A and tdTomato^+^ Basal-B-cell-derived organoids. Scale bars, 50 m. (**Q**) Scatter plot showing the percentage of Ck8^+^ cells in tdTomato^-^ Basal-A and tdTomato^+^ Basal-B-cell-derived organoids. Data are shown as the mean ± SD. Data were analyzed by unpaired two-tailed Student’s t test. \*\**p* < 0.01, ^***^*p* < 0.001, ^****^*p* < 0.0001.

To compare the stemness capacity of Basal-A and Basal-B-cell populations, Markov-chain entropy (MCE) analysis and organoid formation assays were performed. The MCE analysis demonstrated that the Basal-B-cell population had a significantly higher stemness potency score than that of the Basal-A cell population (Fig. 2D). We isolated Basal-A cells (C1rb^low^/Lin^-^/Epcam^high^/CD49f^high^) and Basal-B cells (C1rb^high^/Lin^-^/Epcam^high^/CD49f^high^) from wild-type (WT) B6 mouse prostates by FACS based on the different expression levels of the Basal-B-cell surface marker gene C1rb (Fig. 2, B and E and fig. S5, A to D) and compared their organoid-forming efficiency as previously described (*10*). We found that Basal-B cells exhibited significantly higher organoid-forming capacity than that of Basal-A cells (Fig. 2, F and G). Strikingly, most of the Basal-A-derived organoids formed solid spheres, whereas the Basal-B-derived organoids formed hollow spheres (Fig. 2, F, H and I). Immunofluorescence analysis revealed that compared with Basal-A-derived solid organoids, Basal-B-derived hollow organoids contained more Ck8^+^ luminal cells (Fig. 2J).

To directly trace and characterize Basal-B cells, we generated a genetic lineage tracing model for Nkx3.1-expressing cells by crossing *Nkx3*.*1-CreER* with the *R26-tdTomato* reporter line (fig. S5, E and F). Nkx3.1^+^Trp63^+^Ck5^+^tdTomato^+^ Basal-B cells were found in *Nkx3*.*1-CreER*;*R26-tdTomato* (NT) mouse prostates after Tam treatment (Fig. 2K and fig. S5G to I). To further validate the stemness capacity of Basal-B cells, we isolated tdTomato^+^ Basal-B cells (tdTomato^+^/Lin^-^/Epcam^high^/CD49f^high^) and tdTomato^-^ Basal-A cells (tdTomato^-^/Lin^-^/Epcam^high^/CD49f^high^) from NT mouse prostates by FACS (Fig. 2L) and compared their organoid-forming efficiency and gene expression profiles. Consistently, Basal-B cells had significantly higher stemness ability than that of Basal-A cells and generated hollow organoids that contained more Ck8^+^ luminal cells (Fig. 2, M to Q). Notably, bulk RNA-seq analysis of the isolated Basal-B and Basal-A cells revealed that Basal-B cells were enriched in signaling pathways involved in tissue development and wound healing and infection, suggesting that the Basal-B-cell population has potential stem properties (fig. S5, J and K). MCE analysis using the bulk cell RNA-seq data further validated that Basal-B cells had a significantly higher stemness score than that of Basal-A cells (fig. S5L). Moreover, gene set enrichment analysis (GSEA) revealed that the injury-associated regenerative signature was significantly enriched in Basal-B cells (fig. S5M). These results suggest that this newly identified Basal-B-cell population is a potential prostate basal stem cell population.

### Lineage tracing of Basal-B cells during mouse prostate homeostasis and regeneration

To determine the contribution of Basal-B cells to prostate regeneration *in vivo*, we combined *Trp63-DreER* and *Nkx3*.*1-CreER* with the *R26-TLR* reporter line to generate a *Trp63-DreER*;*Nkx3*.*1-CreER*;*R26-TLR* (TN-TLR) mouse model for lineage tracing of Trp63^+^Nkx3.1^+^ Basal-B cells (ZsGreen^+^tdTomato^+^) during mouse prostate homeostasis and regeneration (Fig. 3A). To further verify the labeling specificity, we crossed the *R26-TLR* reporter line with the inducible *Trp63-DreER* or *Nkx3*.*1-CreER* mouse line, which specifically targets Trp63-expressing basal cells (fig. S6A) or Nkx3.1-expressing cells, respectively (fig. S6B). Two weeks after Tam administration, we collected prostates from three different genotypes: TN-TLR, *Trp63-DreER*;*R26-TLR* (T-TLR) and *Nkx3*.*1-CreER*;*R26-TLR* (N-TLR) (Fig. 3B and fig. S6C). As expected, immunostaining for ZsGreen, tdTomato, Trp63, Nkx3.1 and Ck8 on TN-TLR, T-TLR and N-TLR mouse prostates showed that all ZsGreen^+^ cells were Trp63^+^Ck8^-^ basal cells, and all tdTomato^+^ cells were Nkx3.1-expressing cells (Fig. 3, C and D and fig. S6, D to K). ZsGreen^+^tdTomato^+^Nkx3.1^+^ Basal-B cells were successfully labeled in TN-TLR mouse prostates (Fig. 3, C and D).

**Fig. 3.**
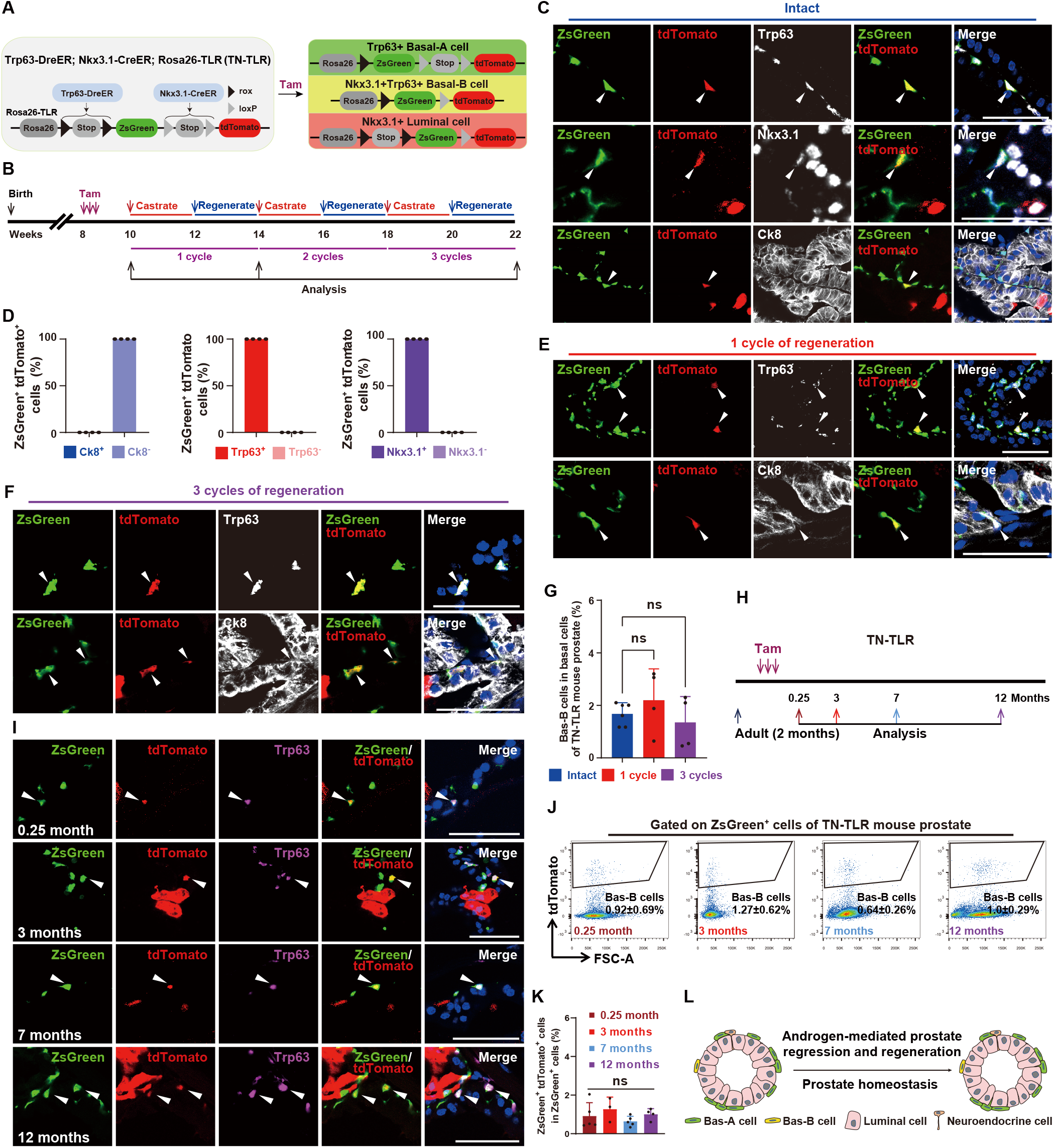
Prostate regeneration and homeostasis are not driven by specific basal stem cell populations. (**A**) Construction strategy of TN-TLR (*Trp63-DreER;Nkx3*.*1-CreER;R26-RSR-ZsGreen-LSL-tdTomato*) mice for Basal-A, Basal-B and Nkx3.1^+^ luminal cell labeling. (**B**) Schedule of Basal-B-cell labeling in TN-TLR mice during prostate regression and regeneration. (**C**) Immunofluorescence staining of Trp63, Nkx3.1, Ck8, ZsGreen and tdTomato in intact TN-TLR mouse prostates. (**D**) Bar graphs showing the percentage of Ck8^+^ (*n* = 0), Ck8^-^ (*n* = 2,004), Trp63^+^ (*n* = 1,557), Trp63^-^ (*n* = 0), Nkx3.1^+^ (*n* = 2,238) and Nkx3.1^-^ (*n* = 0) cells among ZsGreen^+^tdTomato^+^ cells of intact TN-TLR mouse prostates (*n* = 4). (**E** and **F)** Immunofluorescence staining of Trp63, Ck8, ZsGreen and tdTomato in TN-TLR mouse prostates after 1 cycle (E) and 3 cycles (F) of prostate regression and regeneration. (**G**) Bar graph showing the percentage of Basal-B cells (tdTomato^+^ZsGreen^+^) among ZsGreen^+^ basal cells of intact (*n* = 6 mice) and regenerated TN-TLR mouse anterior prostates after 1 cycle (*n* = 4 mice) or 3 cycles (*n* = 4 mice) of regression-regeneration. (**H**) Schedule of Basal-B-cell labeling in TN-TLR mice during long-term prostate homeostasis. (**I**) Immunofluorescence staining of Trp63, ZsGreen and tdTomato in TN-TLR mouse anterior prostates at 4 time points (1 week, 3 months, 7 months and 12 months) after tamoxifen injection. (**J** and **K**) FACS plots (J) and bar graphs (K) showing the percentage of Basal-B cells (tdTomato^+^/ZsGreen^+^) among basal cells (ZsGreen^+^) of TN-TLR mouse anterior prostates at 4 time points (1 week (*n* = 5 mice), 3 months (*n* = 3 mice), 7 months (*n* = 5 mice) and 12 months (*n* = 4 mice)) after tamoxifen injection. (**L**) Schematic diagram suggesting that Basal-B cells do not drive androgen-mediated prostate regeneration or long-term prostate homeostasis. White arrows indicate Basal-B cells (tdTomato^+^ZsGreen^+^). Data are shown as the mean ± SD. Data were analyzed by unpaired two-tailed Student’s t test; ns, nonsignificant.

Next, we performed prostate regression-regeneration assays by castration and testosterone administration cycles in TN-TLR mice (Fig. 3B). ZsGreen^+^tdTomato^+^ Basal-B cells represented 1.68 ± 0.43% of TN-TLR mouse prostate basal cells 14 days after Tam treatment, but the number of ZsGreen^+^tdTomato^+^ labeled Basal-B cells did not significantly change after one cycle (2.2 ± 1.19%) or three cycles (1.35 ± 0.98%) of androgen -mediated prostate regression-regeneration (Fig. 3, E to G). The FACS analysis also showed that the percentage of ZsGreen^+^tdTomato^+^ cells did not significantly change after one or three cycles of prostate regression-regeneration (fig. S7). These results suggest that prostate basal cell regeneration is not specifically driven by Basal-B cells.

Next, we systematically characterized the behavior of basal cells and luminal cells *in vivo* by long-term lineage tracing in ProTracer mice. We analyzed ProTracer mouse prostates at 1 week, 3 months, 7 months and 12 months after Tam induction (fig. S8A). Immunostaining for GFP, Trp63 and Ck8 showed that both Trp63^+^ basal cells and Ck8^+^ luminal cells exhibited extensive cell proliferation during long-term prostate homeostasis (fig. S8, B and C). Quantification of the percentage of GFP^+^ basal and luminal cell at three time windows (months 0.25 to 3, 3 to 7, and 7 to 12) revealed the monthly proliferation rate of the prostate basal and luminal cells, which was 0.015 and 0.032 per month during homeostasis, respectively (fig. S8, D and E). FACS analysis also showed that the percentage of GFP^+^ basal and luminal cells significantly increased during long-term prostate homeostasis (fig. S8, F to I).

To characterize the function of Basal-B cells *in vivo* during homeostasis, we analyzed the basal cell populations in TN-TLR mouse prostates at 1 week, 3 months, 7 months and 12 months after Tam induction (Fig. 3H). Immunostaining for ZsGreen, tdTomato, Trp63 and Ck8 showed that the number of ZsGreen^+^tdTomato^+^ Basal-B cells remained almost constant and that basal cell identity was maintained during long-term prostate homeostasis (Fig. 3I and fig. S9A). FACS analysis also showed that the percentage of ZsGreen^+^tdTomato^+^ cells did not significantly change during long-term prostate homeostasis (Fig. 3, J and K). Immunostaining for Nkx3.1 and Trp63 showed that the percentage of Nkx3.1^+^Trp63^+^ Basal-B cells did not significantly change during long-term prostate homeostasis (fig. S9, B and C). Taken together, both lineage tracing and ProTracer data indicate that adult prostate regeneration and homeostasis are not driven by specific basal stem cell populations (Fig. 3L).

### Lineage tracing of Basal-B cells during mouse prostate inflammation

RNA-seq analysis revealed that Basal-B cells were enriched in signaling pathway of response bacterium infection and injury-associated regenerative signatures (fig. S5, K and M). To investigate the function of Basal-B cells in response to prostate inflammation, we induced prostate inflammation in TN-TLR mouse anterior prostates by *E. coli* infection (Fig. 4A). ZsGreen^+^tdTomato^+^ (Trp63^+^Nkx3.1^+^) Basal-B and ZsGreen^+^ (Trp63^+^Nkx3.1^-^) Basal-A cells were simultaneously traced at after *E. coli* infection. Consistent with previous studies (*16, 40*), immune cell infiltration and epithelial hyperplasia were observed in the inflamed area of TN-TLR mouse anterior prostates 4 weeks after bacterial infection (Fig. 4, B to D). We did not detect any ZsGreen^+^ luminal cells from TN-TLR mouse prostates in the absence of bacterial infection (Fig. 4E). In the inflamed prostates, however, we found ZsGreen^+^ cells that coexpressed the luminal cell marker Ck8 (Fig. 4F), demonstrating that basal cells could reactivate multipotency and generate luminal cells during prostate inflammation. The percentage of ZsGreen^+^tdTomato^+^ Basal-B cells in ZsGreen^+^ basal cells was significantly increased after bacterial infection (from 1.91 ± 0.64% to 6.62 ± 1.24%) (Fig. 4G and H). Moreover, we found that ZsGreen^+^tdTomato^+^ Basal-B cells exhibited approximately fivefold higher basal-to-luminal differentiation efficiency than ZsGreen^+^ Basal-A cells during prostate inflammation (Fig. 4H). In line with this finding, the percentage of Ck8^+^ luminal cells in Basal-B-derived organoids was significantly higher than that in Basal-A-derived organoids (Fig. 2, J and Q). These results demonstrate that Basal-B cells are multipotent prostate stem cells under *in vitro* organoid culture and bacterial infection-induced prostate inflammation conditions (Fig. 4I).

**Fig. 4.**
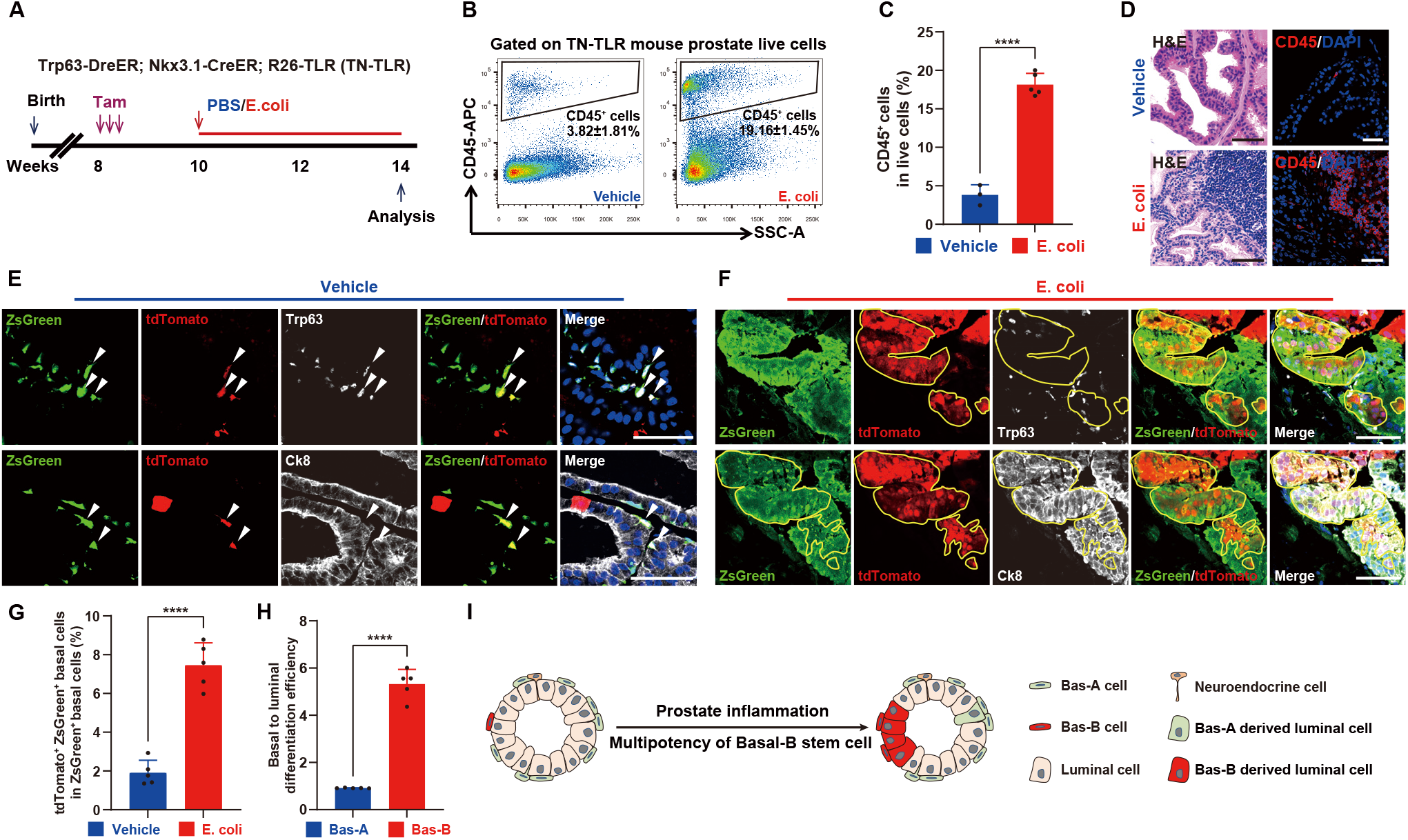
Basal-B cells serve as prostate stem cells in bacteria-induced prostate inflammation. (**A**) Schematic of labeling Basal-B cells and *E. coli* infection in TN-TLR mouse prostates. (**B**) FACS plots showing the percentage of CD45^+^ immune cells in TN-TLR mouse prostates with or without *E. coli* infection. (**C**) Bar plot illustrating the percentage of CD45^+^ immune cells in TN-TLR mouse prostates with or without *E. coli* infection (*n* = 3 mice (Vehicle), *n* = 5 mice (*E. coli* infection)). (**D**) Hematoxylin-eosin staining (H&E) and immunofluorescence staining of CD45 in TN-TLR mouse prostates with or without *E. coli* infection. (**E** and **F**) Immunofluorescence staining of Trp63, Ck8, ZsGreen and tdTomato in TN-TLR mouse prostates with vehicle treatment (E) or *E. coli* infection (F). White arrows indicate ZsGreen^+^tdTomato^+^ Basal-B cells. (**G**) Bar diagram shows the percentage of ZsGreen^+^tdTomato^+^ Basal-B cells in ZsGreen^+^ basal cells in TN-TLR mouse prostates with or without *E. coli* infection (*n* = 5 mice per group). (**H**) Bar plots showing basal-to-luminal cell differentiation efficiency of Basal-A and Basal-B cells in TN-TLR mouse prostates (*n* = 5 mice). (**I**) Schematic diagram suggesting that multipotent Basal-B cells serve as prostate stem cells in bacteria-induced prostate inflammation. Scale bars, 50 m. Data are shown as the mean ± SD. Data were analyzed by unpaired two-tailed Student’s t test; ^****^*p* < 0.0001.

### Basal-B-specific Tracer reveals an increased Basal-B-cell population

To continuously and specifically monitor the Basal-B-cell population, we next modified the TN-TLR model to enable Basal-B-specific recording and tracing. We crossed *Trp63-DreER* with *Nkx3*.*1-xER* mice and with dual recombinase-activated reporter *R26-TLR* mice to generate the Basal-B Tracer mouse model (Fig. 5A). *Trp63-DreER* recombined almost all basal cells after Tam injection (Fig. 3, A to D). Tam-induced DreER-rox recombination excised both rox-ER-rox from *Nkx3*.*1-CrexER* and rox-stop-rox from *R26-TLR*, genetically generating *Nkx3*.*1-Cre* and *R26-rox-ZsGreen-loxp-stop-loxp-tdTomato* genotypes specifically in Trp63-expressing basal cells. Subsequently, upon any event that would increase the number of Basal-B cells, *Nkx3*.*1-Cre* should excise loxP-stop-loxP from *R26-rox-ZsGreen-loxp-stop-loxp-tdTomato*, leading to constitutive ZsGreen and tdTomato expression in Basal-B cells and their daughter cells (Fig. 5A). Immunostaining for ZsGreen, tdTomato, Trp63 and Ck8 on prostate sections from Basal-B Tracer mice revealed that all Zsgreen^+^tdTomato^+^ cells were verified as Nkx3.1^+^Trp63^+^ Basal-B cells; no ZsGreen^+^tdTomato^+^ signal was detected in Ck8^+^ luminal cells two weeks after Tam administration (Fig. 5B). These findings confirm that Basal-B cells were genetically recorded by the Basal-B Tracer.

**Fig. 5.**
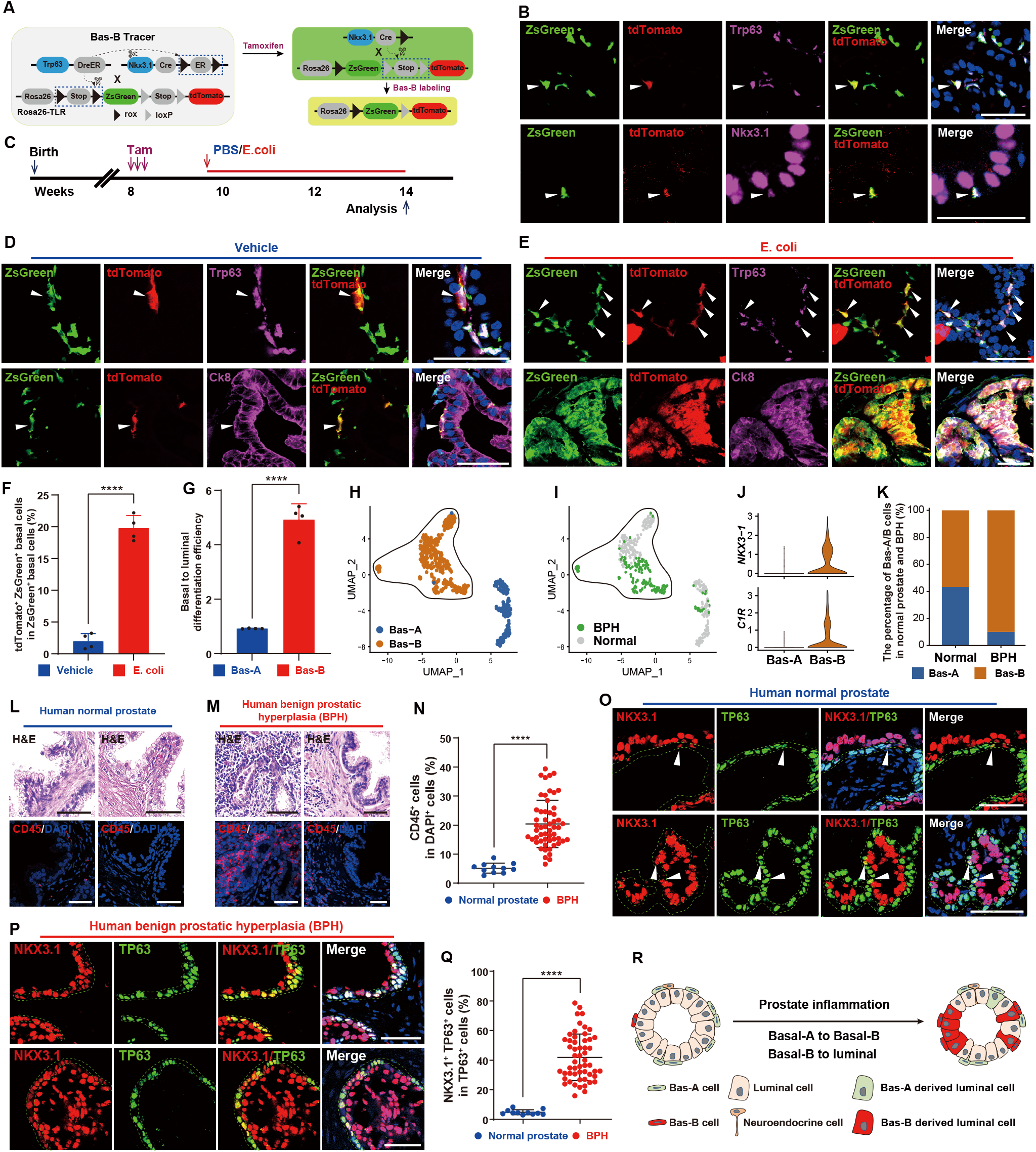
Basal-A-to-Basal-B-to-luminal cell transitions in prostate inflammation and BPH. (**A**) Construction strategy of Basal-B Tracer model (*Trp63-DreER;Nkx3*.*1-CrexER;R26-RSR-ZsGreen-LSL-tdTomato*) mice for enhanced Basal-B-cell tracing. (**B**) Immunofluorescence staining of Trp63, Nkx3.1, ZsGreen and tdTomato in intact Basal-B Tracer mouse prostates. (**C**) Schedule of tracing Basal-B cells and *E. coli* infection in Basal-B tracer mouse prostates. (**D** and **E**) Immunofluorescence staining of Trp63, Ck8, ZsGreen and tdTomato in Basal-B Tracer mouse prostates with vehicle treatment (D) or *E. coli* infection (E). White arrows indicate ZsGreen^+^tdTomato^+^ Basal-B cells. (**F**) Bar graph showing the percentage of ZsGreen^+^tdTomato^+^ Basal-B cells among ZsGreen^+^ basal cells in Basal-B Tracer mouse prostates with or without *E. coli* infection (*n* = 4 mice per group). (**G**) Bar graph showing the efficiency of basal-to-luminal cell transition of Basal-A and Basal-B cells in Basal-B Tracer mouse prostates (*n* = 4 mice per group). (**H**) UMAP plot showing the integrated clustering result of Basal-A and Basal-B cells from human normal prostate and BPH samples. (**I**) UMAP plot showing the Basal-B cell distribution in human normal prostate and BPH samples. (**J**) The gene expression levels of Basal-B markers *NKX3*.*1* and *C1R* in Basal-A and Basal-B cells. (**K**) Cell proportion of Basal-A and Basal-B cells across normal human prostate and BPH samples. (**L** and **M**) H&E and immunofluorescence staining of CD45 in normal human prostate (L) and BPH samples (M). (**N**) Scatter plot showing the percentage of CD45^+^ cells in human normal prostate (*n* = 11) and BPH samples (*n* = 54). (**O** and **P**) Immunofluorescence staining of NKX3.1 and TP63 in human normal prostate (O) and BPH samples (P). White arrows indicate TP63^+^NKX3.1^+^ Basal-B cells in the normal prostate samples. (**Q**) Scatter plot showing the percentage of TP63^+^NKX3.1^+^ Basal-B cells among TP63^+^ basal cells in human normal prostate (*n* = 11) and BPH samples (*n* = 54). (**R**) Schematic diagram suggests Basal-A-to-Basal-B-to-luminal cell transitions in prostate inflammation and BPH. Scale bars, 50 m. Data are shown as the mean ± SD. Data were analyzed by unpaired two-tailed Student’s t test; ^****^*p* < 0.0001.

To continuously record the Basal-B cell population in response to prostate inflammation, we induced prostate inflammation in Basal-B tracer mouse prostates (Fig. 5C). Consistently, we did not observe significant cell number changes in ZsGreen^+^tdTomato^+^Trp63^+^ Basal-B cells and did not detect any ZsGreen^+^tdTomato^+^ luminal cells from Basal-B Tracer mouse prostates in the absence of bacterial infection (Fig. 5D). The percentage of ZsGreen^+^tdTomato^+^ Basal-B cells and their daughter cells were significantly increased in ZsGreen^+^ basal cells after bacterial infection (from 1.99 ± 1.2% to 19.78 ± 2.01%) (Fig. 5, E and F). We further confirmed that ZsGreen^+^tdTomato^+^ Basal-B cells had a significantly higher basal-to-luminal differentiation efficiency than that of ZsGreen^+^ Basal-A cells during prostate inflammation (Fig. 5, E and G).

### Basal-B cell population in human prostate samples

Asymptomatic prostatic inflammation is very common in adult men, including prostate tissues removed for the treatment of both prostate cancer and benign prostatic hyperplasia (BPH) (*17*). To investigate the presence of NKX3.1-expressing Basal-B cells in prostates from normal individuals and individuals with BPH, we integrated the published human prostate sc-RNA-seq datasets (*27, 29*), which include data from both human normal prostate and BPH samples, and specifically analyzed the basal cell populations (fig. S10). We do find an h-Basal-B cell cluster in human normal prostates and BPH samples, with high *TP63, CK5, NKX3*.*1* and *C1R* expression (Fig. 5, H to J and fig. S10). Strikingly, the percentage of Basal-B cells was dramatically increased in BPH samples compared with normal prostate samples (Fig. 5K). These results further confirm the existence of the Basal-B cell population in the prostates of normal human individuals and individuals with BPH.

To validate the Basal-B cell population in human prostates, we investigated Basal-B cell marker gene expression levels and the inflammation status in human normal prostate and BPH samples. Hematoxylin and eosin (H&E) staining and CD45 immunofluorescence staining showed that CD45^+^ immune cell infiltration was significantly increased in BPH samples compared with human normal prostate samples (Fig. 5, L to N). Strikingly, immunostaining for NKX3.1 and TP63 showed that the percentage of NKX3.1^+^TP63^+^ Basal-B cells was significantly increased in human BPH samples compared with normal prostate samples (Fig. 5, O to Q). These results demonstrate that the Basal-B cell population represents the dominant basal cell type in BPH, suggesting the potential prostate basal cell lineage plasticity of the Basal-A-to-Basal-B transition during prostatic inflammation (Fig. 5R).

### Basal cell lineage plasticity in basal cell-initiated prostate intraepithelial neoplasia (PIN)

To investigate the functions of the whole basal cell population *in vivo* during prostate cancer initiation, we generated a basal cell-specific lineage tracing model by knocking in the *CreER* cassette under control of the *Ck5* promoter. We crossed *Ck5-CreER* mice with *R26-tdTomato* and *Pten*^*flox/flox*^ mice to generate *Ck5-CreER*;*R26-tdTomato*;*Pten*^*flox/flox*^ (C5TP) mice (Fig. 6A). Quantification of the percentage of tdTomato^+^Ck5^+^ basal cells after 2 days of Tam treatment revealed that the basal cell labeling efficiency was approximately 96.12 ± 2.4% (Fig. 6, B and C and fig. S11, A and B). Fate mapping of tdTomato^+^ basal cells at five time points (2 days, 1 week, 2 weeks, 3 weeks, and 4 weeks after Tam treatment) revealed that the percentage of tdTomato^+^Nkx3.1^+^Ck5^+^Pten^-^ Basal-B cells was significantly increased in tdTomato^+^ basal cells (from 8.72 ± 1.61% to 89.06 ± 1.88%) at the time window of 2 days to 1 week after Tam treatment (Fig. 6, C to H), suggesting extensive Basal-A-to-Basal-B transition after Pten deletion in the prostate basal cell population. The percentage of tdTomato^+^Ck8^+^Pten^-^ luminal cells gradually emerged (0.33 ± 0. 14%) at 1 week after Tam treatment and significantly increased (3.48 ± 0. 47%) at 2 weeks after Tam treatment (Fig. 6, C to I), suggesting potential Basal-B-to-luminal lineage plasticity after Pten deletion in the prostate basal cell population.

**Fig. 6.**
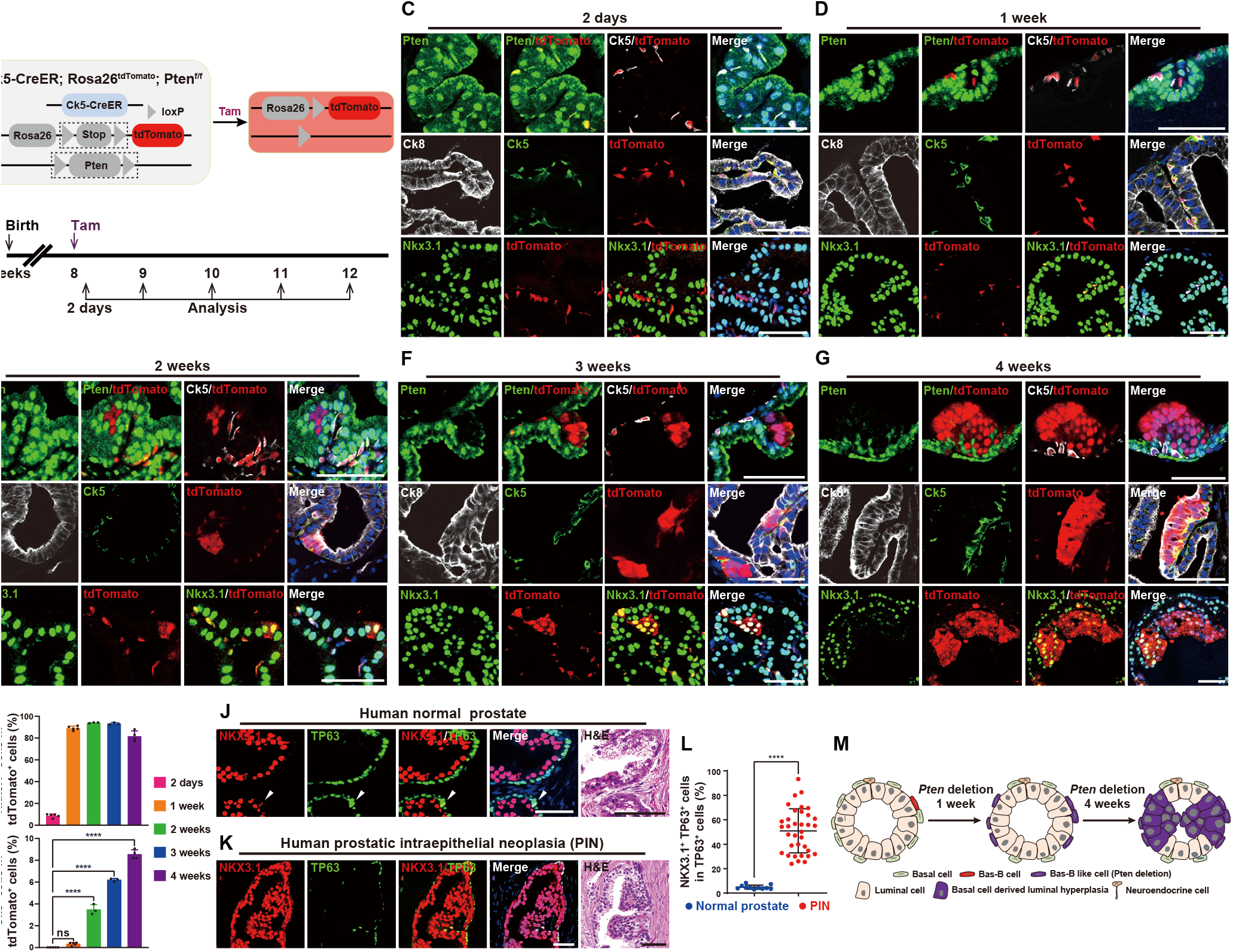
Basal cell lineage plasticity in mouse basal cell-derived prostate hyperplasia and human PIN lesions. (**A**) Construction strategy of *Ck5-CreER;R26-LSL-tdTomato;Pten*^*flox/flox*^ (K5CP) mice for basal cell tracing during prostate cancer initiation induced by Pten deletion. (**B**) Schedule of basal cell tracing in K5CP mouse prostates. (**C**-**G**) Immunofluorescence staining of Pten, Ck5, Nkx3.1, Ck8 and tdTomato in K5CP mouse prostates at 2 days (C), 1 week (D), 2 weeks (E), 3 weeks (F) and 4 weeks (G) after tamoxifen treatment. (**H** and **I**) Bar graphs showing the percentage of Nkx3.1^+^ cells (H) or Ck8^+^ cells (I) among tdTomato^+^ cells in K5CP mouse prostates at 2 days (n = 5), 1 week (n = 5), 2 weeks (n = 3), 3 weeks (n = 3) and 4 weeks (n = 4) after tamoxifen treatment. (**J** and **K**) H&E staining and immunofluorescence staining of NKX3.1 and TP63 in human normal prostate (J) and PIN samples (K). (**L**) Scatter plot showing the percentage of TP63^+^NKX3.1^+^ Basal-B cells among TP63^+^ basal cells in human normal prostate (*n* = 11) and PIN samples (*n* = 35). (**M**) Schematic diagram suggesting the potential Basal-A-to-Basal-B-to-luminal cell lineage plasticity during basal cell-induced prostate cancer initiation. Scale bars, 50 m. Data are shown as the mean ± SD. Data were analyzed by one-way ANOVA; ^****^*p* < 0.0001 and ns, nonsignificant.

To validate the potential existence of NKX3.1-expressing Basal-B cells under precancerous conditions, we performed NKX3.1 and TP63 immunofluorescence staining in human PIN samples (Fig. 6, J and K). Interestingly, the percentage of the NKX3.1^+^TP63^+^ Basal-B cell population was significantly increased in human PIN samples compared with normal prostate samples (Fig. 6L), indicating potential Basal-A-to-Basal-B lineage plasticity during human prostate cancer initiation.

Taken together, the results of mouse basal cell fate mapping during prostate cancer initiation and the increased Basal-B cell population in human PIN lesions indicate that Basal-A-to-Basal-B lineage plasticity may underlie basal cell-initiated prostate cancer (Fig. 6M).

## Discussion

This work has identified Basal-B cells as an intermediate basal cell population that exhibits greater lineage plasticity under prostate inflammation and cancer initiation conditions. Using a genetic proliferation cell tracing system, enhanced lineage tracing strategies, and superefficient basal cell labeling tools, we uncovered that Nkx3.1-expressing intermediate Basal-B cells have a greater tendency to generate luminal cells *in vitro* and under bacteria-induced prostate inflammation *in vivo*. Strikingly, tremendous Basal-A-to-Basal-B-to-luminal cell transitions have been identified in basal cell-initiated PIN lesions. NKX3.1-expressing Basal-B cells are present in human normal prostates and their abundance is significantly increased in BPH and PIN samples. In addition, our data challenge the intuitive notion that basal cells with low AR expression levels are largely unaffected by androgen deprivation therapy (*19-21*).

Whether there are specific adult stem cell population response for prostate homeostasis, regeneration and cancer initiation is controversial. Using individual markers and lineage tracing techniques, previous studies have proven the existence of adult basal stem cells and luminal stem cells (*21, 33-35*). The potential nonspecific labeling is the main caveat of these genetic lineage-tracing methods because Cre recombinase are supposed to be activated under the control of ‘cell-specific’ markers (*41*). The evidence in support of Ck5^+^ adult basal stem cells generating luminal cells is based on the idea that Ck5 is not expressed in luminal cells (*34, 35*). However, the contribution of adult basal and luminal stem cells requires independent reassessments under different physiological and pathological conditions. In this study, we generated a dual-recombinase-mediated genetic system to label basal, luminal and intermediate cells and simultaneously monitored their cell fates during adult prostate homeostasis, regeneration, inflammation and cancer initiation. In our system, most of the basal cells were labeled with ZsGreen (>93%) in a TN-TLR model, which is more efficient than any other inducible basal cell tracing strategy that has been published. In line with recent works (*13-15*), our results against that adult basal cells contain multipotent stem cell populations during adult prostate homeostasis and androgen-mediated prostate regeneration (*34, 35*). Notably, NKX3.1 was defined as a marker gene of Basal-B cells in this study. Although more detailed mechanistic studies are needed, our study reinforced the basic role of NKX3.1 in regulating prostate cell lineage plasticity.

Intermediate cells have been identified in human prostates with chronic inflammation (*17, 22*). Ck14-expressing basal cells were proposed as the cell origin of these intermediate cells (*16*). Until now, it has not been possible to distinguish the intermediate cell population among Ck14-expressing basal cells because the defined cell markers were unknown and the necessary strains to fate trace intermediate cells were not available. By utilizing unbiased scRNA-seq technology and enhanced lineage tracing strategies that use dual recombinases, we showed that Nkx3.1 and Trp63 double-positive Basal-B cells contribute to the generation of intermediate cells and luminal cells and provided genetic evidence of Basal-A-to-Basal-B-to-luminal transitions under bacteria-induced prostate inflammation. Similar to the mouse model, Basal-B cells were significantly increased in human BPH and PIN samples compared with human normal prostate samples. The identification of such intermediate cells that preferentially exhibit more lineage plasticity under prostatic inflammation and cancer initiation has implications for understanding the cellular basis of prostate cancer.

## Supporting information

Supplementary Materials Description

## Acknowledgments

We thank Baojin Wu and Guoyuan Chen for their help with animal husbandry and Wei Bian for providing technical help at the CEMCS (SIBCB) Core Facility. We would like to thank the Genome Tagging Project (GTP) Center, CEMCS (SIBCB), and CAS for technical support (Shanghai, China).

## Author contributions

D.G. conceived the project. W.X.G., B.Z. and D.G. designed the experiments. W.X.G. and X.Y.Z. performed most of the experiments. H.J.Z., K.L., S.M.W., Y.Y.P., Y.J. and C.Y. helped with the experiments and provided technical support. P.F.S. and C.L. provided human prostate samples. L.L. and L.N.C. performed computational and statistical analyses. B.Z. and K.L. provided the ProTracer and dual recombinases system. L.N.C., B.Z. and D.G. oversaw the project.

## Data and materials availability

All raw sequencing data in this study have been uploaded to the National Omics Data Encyclopedia (NODE; https://www.biosino.org/node/index) with the accession number OEP003997, https://www.biosino.org/node/review/detail/OEV000439?code=4XXKPANR. Previously published scRNA-seq reanalyzed in our manuscript are available under accession codes OEP000825 (PMID: 32807988) and GSE150692 (PMID: 32915138). The custom code used is available at GitHub (https://github.com/LinLi-0909/Prostate_Basal).

## Supplementary Materials

Materials and Methods Figs. S1 to S11

Tables S1 to S6

References (*1-17*)

