## Supplementary Materials Description for "Intermediate basal cell population in prostate homeostasis and cancer initiation": Supplementary Materials.pdf

**This PDF file includes:**

Materials and Methods

Figs. S1 to S11

Tables S1 and S2

References (*1-17*)

**Other Supplementary Materials for this manuscript include the following:**

Tables S3 to S6

### Materials and Methods

#### Mice

All mouse studies were conducted under the guidelines of the Animal Care and Use Committee at the Center for Excellence in Molecular Cell Science, Chinese Academy of Sciences (CAS). All mice were genotyped by genomic PCR, and the specific primers for each mouse line are listed in Supplementary Table 1.

The mouse lines *Rosa26-loxp-stop-loxp-tdTomato*, *Pten*$^{flox/flox}$, *Ck8-CreER*, *Rosa26-TLR*, *Rosa26-DreER*, *Rosa26-loxp-stop-loxp-GFP*, *Ki67-CrexER*, and *Trp63-DreER* were reported in a previous study (*1-5*). In brief, the *Rosa26-loxp-stop-loxp-tdTomato* mouse line was generated by knocking the CAG-loxP-stop-loxP-tdTomato-WPRE-pA-AttB-Pr-FRT-Neo-pA-Attp cassette into the intron between exons 1 and 2 of the *Rosa26* gene locus. The *Pten*$^{flox/flox}$ mouse line was generated in which exons 4 and 5 of the *Pten* gene are flanked by loxP sites. For generating *Ck8-CreER*, a BAC clone RP23-234D2 was first purchased from the BACPAC Resource Center which comprises the full-length *CK8* gene and 60 kb of 5′ and 100 kb of 3′ flanking sequences. And the transgene unit, contains Cre-ERT2 cDNA and an IRES–GFP expression cassette, was inserted at the translational initiation codon of the Ck8 gene of BAC clone RP23-234D2. The *R26-TLR* mouse line was generated by knocking the CAG-rox-stop-rox-ZsGreen-WPRE-pA-Frt-Neo-Frt sequence and the insulator-CAG-loxP-stop-loxP-tdTomato-WPRE-pA sequence into exons 1 and 2 of the *Rosa26* gene locus using homologous recombination, respectively. For generating the *Rosa26-DreER* mouse line, the target vector containing pCAG-DreERT2-BGHpA-Neo-DTA sequence was inserted into the *Rosa26* locus to generate a *Rosa26-DreER* allele. For generating the *Rosa26-loxp-stop-loxp-GFP* mouse line, the target vector containing CAG-loxP-stop-loxP-EGFP-WPRE sequence was inserted into the exon 2 of *Rosa26* locus to generate a Rosa26-loxp-stop-loxp-GFP allele. For generating *Ki67-CrexER* (*Ki67-Cre-rox-ER-rox*), the translational stop codon of the *Ki67* gene locus was inserted into a cDNA encoding CrexER recombinase by insertion of two rox sequences flanking ERT2. The *Trp63-DreER* mouse line was generated by knocking the DreERT2-WPRE-polyA sequence into the exon 4 of the *Trp63* gene locus.

*Nkx3.1-CreER* and *Ck5-CreER* were purchased from Shanghai Model Organisms Center. In brief, the *Nkx3.1-CreER* mouse line was generated by knocking the IRES-CreERT2 sequence into the 3' UTR site of Nkx*3.1* gene locus. The *Ck5-CreER* mouse line was generated by targeting 2A-CreERT2 cassette into the translational stop codon of *Ck5* gene locus through homologous recombination using CRISPR/Cas9.

The new mouse line *Nkx3.1-CrexER* was generated by CRIPR-Cas9 technology. In brief, the

*Nkx3.1-CrexER* mouse line was generated by knocking the 2A-Cre-rox-ERT2-rox sequence into the 3' UTR site of *Nkx3.1* gene locus.

***In vivo* prostate castration and regeneration**

Male mice were castrated at the age of 10 weeks using standard procedures. Testosterone pellets (12.5 mg per pellet, 90-d release, Innovative Research of America) were implanted subcutaneously after two weeks of castration to restore the serum testosterone level and stimulate prostate regeneration. The testosterone pellets were removed to reinduce prostate regression 2 weeks after prostate regeneration. Testosterone pellets were subcutaneously transplanted again to reinduce prostate regeneration. After 2 weeks of this regeneration, the mice had undergone two cycles of androgen-mediated prostate regression-regeneration.

**The *E. coli* strain and *E. coli*-mediated prostatic injury model**

A single *E. coli* (DH5α) strain was inoculated into 5 mL of Luria broth (LB) medium at an ultraclean workbench and cultured for 12 hours at 37°C. Then, the *E. coli* suspension was passaged into 50 mL LB medium at a 1:10 ratio at 37°C. The *E. coli* suspension was adjusted to an A600 value of 0.85 (bacterial concentration equal to $4.5 \times 10^8$ cfu/mL), washed and concentrated with sterilized PBS to a final concentration of $1 \times 10^{10}$ cfu/mL. Experimental mice were anesthetized by 3% isoflurane in the anesthesia box and transferred to a 37°C thermostatic mat. Then, anesthesia was maintained with 2% isoflurane and oxygen gas. After removing the abdominal fur and disinfecting the skin with an iodophor-soaked cotton ball, a lower abdominal cutaneous and muscular incision was performed to expose the prostate. Twenty-five microliters of the $1 \times 10^{10}$ cfu/mL *E. coli* suspension was injected into each anterior prostate. The cutaneous and muscular incision was sutured with absorbable sutures and suture nails. After the surgical operation, the mice were moved to a warming plate and transferred into mouse cages.

**Prostate harvesting and fluorescence-activated cell sorting (FACS)**

The mice were sacrificed using $CO_2$. Prostate tissues were removed and minced under a stereomicroscope. Then, the prostate tissues were incubated with 1 mL 1× collagenase/hyaluronidase (STEMCELL, no. 07912) with 10 μM Y27632 at 37°C with gentle agitation for 30 min. Following centrifugation at 800 rpm for 1 min and removal of the supernatant, the dissociated tissue samples were further digested in TrypLE (Gibco, no. 12605-028) with 10 μM Y27632 for 10 min at 37°C with gentle shaking. Following quenching of TrypLE using DMEM with 10% FBS, the dissociated cells were stained with antibodies for 30 min on ice (Supplementary Table 2) and washed with FACS buffer (1x PBS + primocin + 10 mM HEPES + 1% BSA). The prostate cell samples were passed through a 45 μm filter. Cells were sorted using a FACS Aria III (BD).

**Prostate organoid formation assay**

The organoid formation assay was carried out as previously reported (*6, 7*). The sorted prostate cells from experimental mice were resuspended using mouse prostate organoid medium, mixed with an equal volume of Matrigel (Corning 354234) at a final concentration of 250 or 500 cells per 50 μL droplet, and placed into a 24-well plate. The 24-well plate was incubated in a cell incubator (5% $CO_2$, 37°C) for 0.5 hours. The Matrigel was covered with 1 mL mouse prostate organoid medium, the medium was replenished after 4 days of culture, and the organoids were imaged after 7 days of culture using Cytation 5. Organoid measurements were conducted using Gen 5 Image prime.

**Bulk RNA library construction and sequencing**

Ten to twenty thousand Basal-A and Basal-B cells were sorted for each RNA library sample, and RNA isolation and library construction were performed using the manufacturer's instructions. An RNeasy Mini Kit (QIAGEN, no. 74106) was used for RNA isolation, and an mRNA-seq V3 Library Prep Kit (Vazyme, no. NR611) was used for library construction. The samples were sequenced on a NovaSeq6000 (Illumina).

**Immunostaining and hematoxylin-eosin staining assays**

Mice were sacrificed using $CO_2$. Prostate tissues were dissected from sacrificed mice and fixed in 4% paraformaldehyde at 4°C for 2 hours. The tissues were washed with precooled PBS three times and dehydrated by precooled 30% sucrose in PBS overnight at 4°C. Dehydrated tissues were transferred into 30% OCT in 30% sucrose for 1 hour at 4°C. Tissues were embedded in OCT and stored at -80°C. Frozen samples were cut into 10 μm sections and stored at -20°C. Tissue paraffin embedding was conducted using a standard protocol. Paraffin-embedded samples were cut into 5 μm sections and stored at room temperature. Paraffin-embedded sections were rehydrated and underwent antigen retrieval in sodium citrate buffer (pH 6.0) under boiling water bath conditions before immunostaining. Sections were permeabilized in 1× PBS containing 0.1% Triton X-100 and blocked with 5% goat serum in 1× PBS containing 0.1% Triton X-100. Sections were then incubated with primary antibodies in 5% goat serum at room temperature for 2 hours or overnight at 4°C. The sections were washed with PBST (0.05% Tween 20 in 1× PBS) three times before incubation with secondary antibodies in 5%

goat serum at room temperature in darkness for 1 hour. The sections were washed with PBST three times, stained for nuclei with DAPI (Sigma) and mounted with mounting medium. For weak signals, tyramide signal amplification (TSA) was performed following the manufacturer's instructions. The antibodies are listed in Supplementary Table 2.

For hematoxylin-eosin staining, paraffin-embedded sections were stained with hematoxylin for 90 seconds after sections were rehydrated. Sections were washed with running tap water for 5 min. The cells were dehydrated with 1% hydrochloric acid in ethyl alcohol for 2 seconds and then washed with running tap water for 5 min. Sections were stained with eosin for 7 seconds, dehydrated and mounted with neutral resin.

**Preprocessing of bulk RNA-seq data**

We mapped paired-end reads to the GRCm38 mouse transcriptome using hisat2 (v2.1.0) (*8*) with a more than 90% average overall alignment rate. Read counts of RNA-seq were calculated by featureCounts (V1.6.0) (*9*), and a normalized expression matrix was obtained using the fragments per kilobase of exon per million mapped fragments (FPKM) function in the R package Deseq2 (*10*).

**Preprocessing of single-cell data**

**(1) Cluster analysis of basal cells in the mouse prostate atlas**

To analyze the distribution of basal cells in the mouse normal prostate atlas, we extracted basal cell data from a published dataset (*1*) and performed clustering analysis using Seurat 3 (*11*). We excluded cells with less than 200 genes and more than 5,000 genes expressed, and a total of 746 basal cells were used for further analysis. Principal component analysis (PCA) and T-Distributed Stochastic Neighbor Embedding (t-SNE) were performed using 'RunPCA' and 'RunTSNE' with the parameter settings at dims = 1:10. Then, we applied FindClusters to perform clustering analysis with resolution = 0.16, and two distinct clusters were identified for basal cells.

**(2) Cluster analysis of basal cells in the Human Prostate Atlas**

To more accurately identify basal cells in normal human prostate samples and BPH samples, we performed a cluster analysis on epithelial cells from two published human prostate atlas datasets (*1, 12*). For the normal human prostate atlas dataset, we selected epithelial cells with more than 500 genes. Cells with mitochondrial gene percentages greater than 15% were removed, which resulted in 18,497 features across 9,730 cells. We normalized the data with the scale factor setting as 10,000. A total of

1,000 highly variable genes were selected by 'FindVariableFeatures' with the "vst" selection method. PCA and UMAP were performed based on these highly variable genes and the first 6 significant PCs. Epithelial cells of the human prostate were grouped into 3 clusters (resolution = 0.08). For the BPH dataset, we selected epithelial cells and log-transformed the data (scale factor = 1,000,000). A total of 1,000 highly variable genes were selected by 'FindVariableFeatures' with the "vst" selection method. We further performed PCA, and the top 10 PCs were applied to perform UMAP. Finally, we identified 3 clusters for epithelial cells from the BPH dataset (resolution = 0.04).

**Integration of basal cells from normal human prostate and BPH samples**

To compare the distribution of basal cells in BPH and normal prostate samples, we integrated basal cells from the BPH dataset (*12*) and normal human prostate dataset (*1*). To avoid the large difference in cell numbers affecting the integration analysis, we randomly sampled 300 basal cells from the normal prostate scRNA-seq dataset. Then, basal cells from the BPH and normal prostate datasets were integrated by Harmony (*13*). We first deleted cells with mitochondrial gene percentages greater than 10%. A total of 1,200 variable genes were selected by the function 'FindVariableFeatures'. We further use the RunHarmony function to integrate the two datasets. The top 10 PCs were used for dimension reduction and clustering (resolution = 0.03). Finally, basal cells were grouped into 2 distinct clusters.

**Markov-chain entropy (MCE) analysis for basal cells in the mouse prostate single-cell atlas**

MCE (*14*) values were computed to estimate the differentiation potency or potential of basal cells (Basal-A and Basal-B). We first normalized count data with the 'NormalizeData' function (scale factor = 10,000). Furthermore, we added an offset value of 0.1 to the normalized data and log2-transformed the data. Then, we mapped mouse gene symbols to human homologs using Metascape (*15*) and mapped them to human Entrez gene IDs. Then, the MCE values of basal cells were obtained. We found that the MCE values of Basal-B cells were significantly higher than the MCE values of Basal-A cells (*P* value = $5.8\times10^{-20}$, t test).

**Gene set enrichment analysis of basal cell bulk RNA-seq**

We first excluded genes with a mean expression level less than 1 based on FPKM data from bulk RNA-seq. Then, we mapped mouse gene symbols to human homologs. Furthermore, we identified significantly enriched pathways using gene set enrichment analysis (GSEA) (*16*) based on the normalized data, in which the reference dataset was set as Gene Ontology (GO) biological processes.

To identify the injury regenerative function of basal cells, we applied normalized data to perform

GSEA based on the injury-associated regenerative signature from intestinal organoids (*17*).

**Quantification and statistics**

Significance was assessed by an unpaired two-tailed Student's t test. The organoid formation data were acquired from at least three independent experiments. At least three mice were analyzed for mouse prostate regression-regeneration, prostate homeostasis and *E. coli*-induced prostate inflammation assays. A minimum of 10 sections were analyzed in the immunofluorescence analysis of Trp63, Ck5, Ck8, Nkx3.1, tdTomato and ZsGreen.

Fig. S1

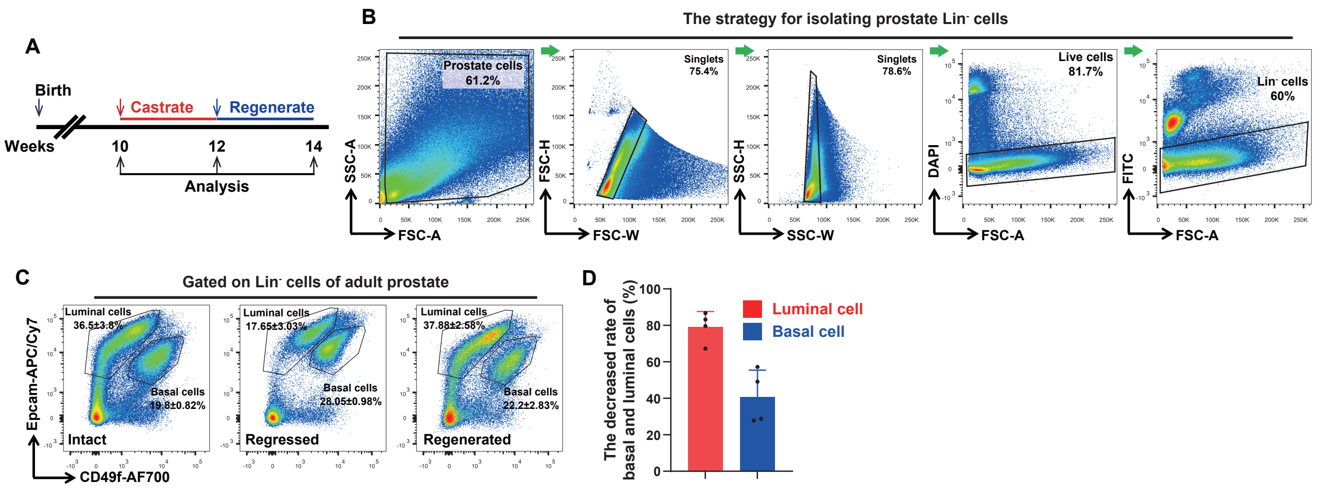

**Fig. S1. Basal cell loss and proliferation during prostate regression and regeneration.** (**A**) Schedule of analyzing total prostate basal and luminal cell numbers during prostate regression and regeneration in wild-type (WT) adult mouse prostates. (**B**) The sorting strategy for isolating prostate Lin⁻ cells by FACS. (**C**) FACS analysis showing the percentage of basal and luminal cells in Lin⁻ cells of intact, regressed and regenerated WT mouse prostates. (**D**) Bar graph showing a decreased percentage of basal and luminal cells in regressed prostates compared to intact prostates ($n$ = 4 mice per group). Data are shown as the mean ± SD.

Fig. S2

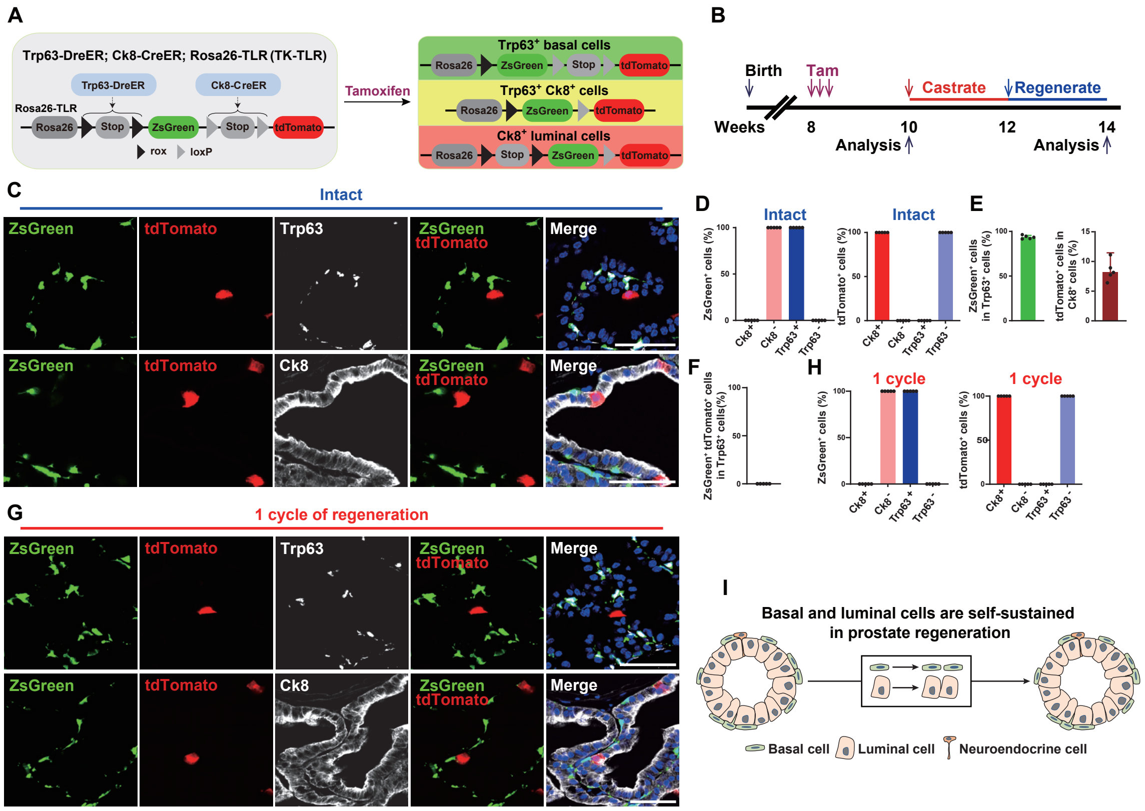

**Fig. S2. Basal and luminal cells are self-sustained during androgen-mediated prostate regression and regeneration.** (**A**) Construction strategy of TK-TLR (*Trp63-DreER;Ck8-CreER;R26-TLR*) mice for Trp63$^+$ basal cell, Ck8$^+$ luminal cell and Trp63$^+$Ck8$^+$ intermediate cell tracing. (**B**) Schedule of basal, luminal and intermediate cell labeling in TK-TLR mice during prostate regression and regeneration. (**C**) Immunofluorescence staining of Trp63, Ck8, ZsGreen and tdTomato in intact TK-TLR mouse prostates. (**D**) Percentage of Ck8$^+$ cells ($n$ = 0), Ck8$^-$ cells ($n$ = 16,479), Trp63$^+$ cells ($n$ = 16,699), and Trp63$^-$ cells ($n$ = 0) among ZsGreen$^+$ cells (left panel); percentage of Ck8$^+$ cells ($n$ = 15,044), Ck8$^-$ cells ($n$ = 0), Trp63$^+$ cells ($n$ = 0), and Trp63$^-$ cells ($n$ = 17,958) among tdTomato$^+$ cells (right panel) of intact TK-TLR mouse prostates ($n$ = 5). (**E**) Percentage of ZsGreen$^+$ cells among Trp63$^+$ basal cells ($n$ = 7,197) (left panel) or tdTomato$^+$ cells among Ck8$^+$ luminal cells ($n$ = 34,350) (right panel) in TK-TLR mouse prostates (n=5). (**F**) Percentage of ZsGreen$^+$tdTomato$^+$ cells ($n$ = 0) in Trp63$^+$ basal cells ($n$ = 24,240) ($n$ = 5 mice). (**G**) Immunofluorescence staining of Trp63, Ck8, ZsGreen and tdTomato in TK-TLR mouse prostates after one cycle of prostate regression-regeneration. (**H**) Percentage of Ck8$^+$ cells ($n$ = 0), Ck8$^-$ cells ($n$ = 22,577), Trp63$^+$ cells ($n$ = 17,958), and Trp63$^-$ cells ($n$ = 0) among ZsGreen$^+$ cells (left panel); Ck8$^+$ cells ($n$ = 18,657), Ck8$^-$ cells ($n$ = 0), Trp63$^+$ cells ($n$ = 0), and Trp63$^-$ cells ($n$ = 16,821) among tdTomato$^+$ cells (right panel) in TK-TLR mouse prostates ($n$ = 5) after one cycle of prostate regression-regeneration. (**I**) Schematic diagram suggesting that basal and luminal cells are self-sustained during androgen-mediated prostate regression and regeneration. Scale bars, 50 µm. Data are shown as the mean ± SD.

Fig. S3

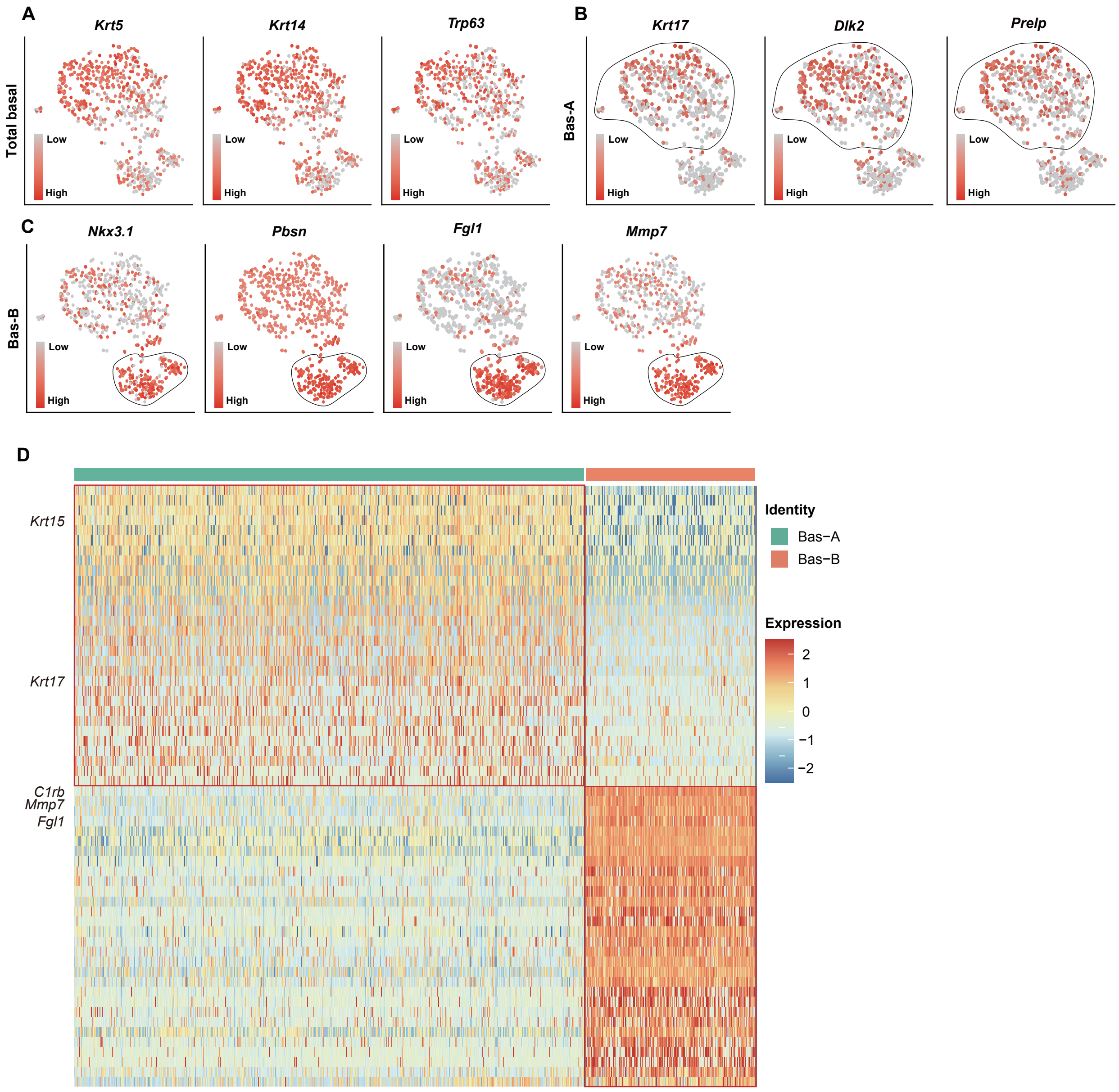

184 **Fig. S3. Characterization of basal cell subtypes in adult mouse prostates.** (**A** to **C**) t-SNE plots
185 show the expression levels of basal signature genes: *Krt5*, *Krt14* and *Trp63,* **(A)** Basal-A signature
186 genes: *Krt17*, *Dlk2* and *Prelp,* **(B)** Basal-B signature genes: *Nkx3.1*, *Pbsn*, *Fgl1*, *Mmp7* and *C1rb,* **(C)**
187 across basal cells. (**D**) Heatmap shows the expression levels of the top 30 signature genes across basal
188 cells.

Fig. S4

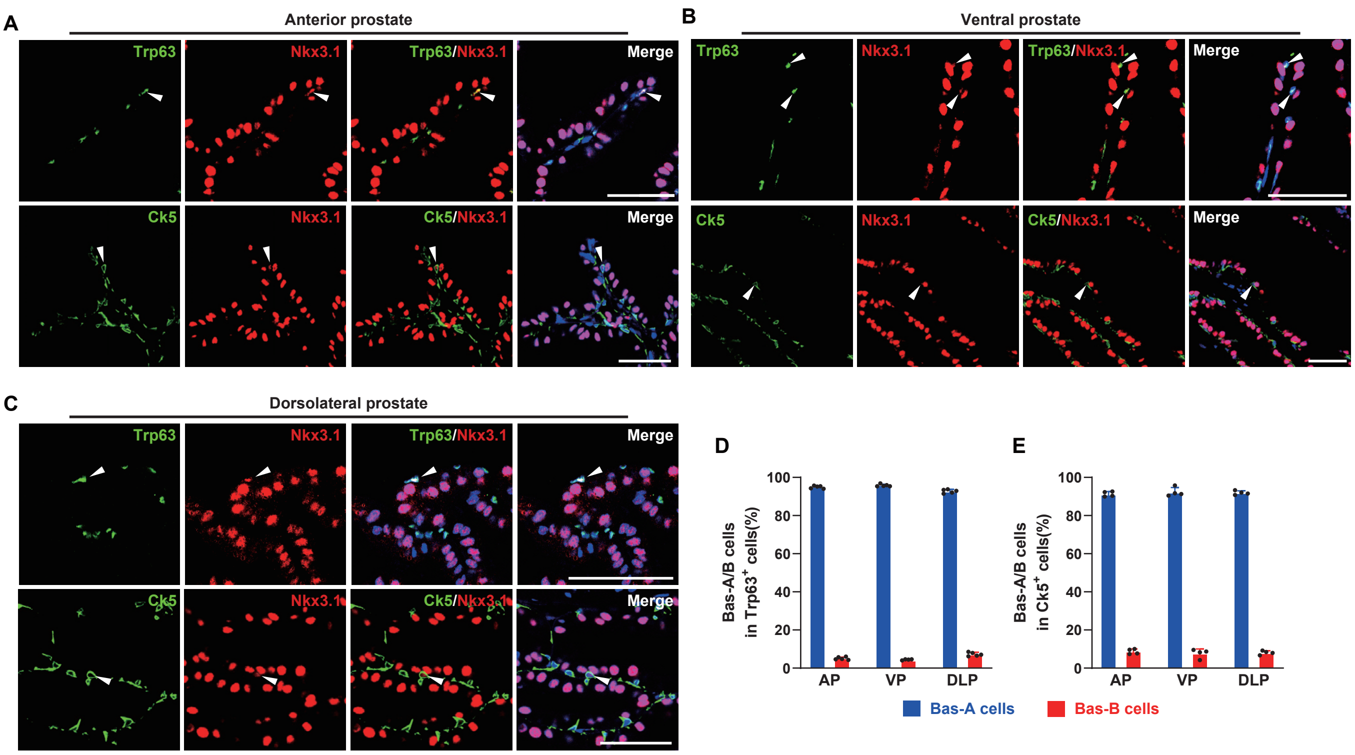

**Fig. S4. Basal-A and Basal-B cells have been identified in adult WT mouse prostates.** (**A** to **C**) Immunofluorescence staining of Trp63, Nkx3.1 and Ck5 in the WT mouse prostate anterior lobe (A), ventral lobe (B) and dorsal-lateral lobe (C). (**D** and **E**) Percentage of Basal-A and Basal-B cells among Trp63$^{+}$ basal cells of the anterior lobe ($n$ = 34,026 cells), ventral lobe ($n$ = 3,360 cells), and dorsal-lateral lobe ($n$ = 3,578 cells) of the prostate in WT mice ($n$ = 5) (D) and Ck5$^{+}$ basal cells of the anterior lobe ($n$ = 7,749 cells), ventral lobe ($n$ = 5,080 cells), and dorsal-lateral lobe ($n$ = 2,520 cells) of the prostate in WT mice ($n$ = 4) (E). Scale bars, 50 µm. White arrows indicate Nkx3.1$^{+}$ Basal-B cells. Data are shown as the mean ± SD.

Fig. S5

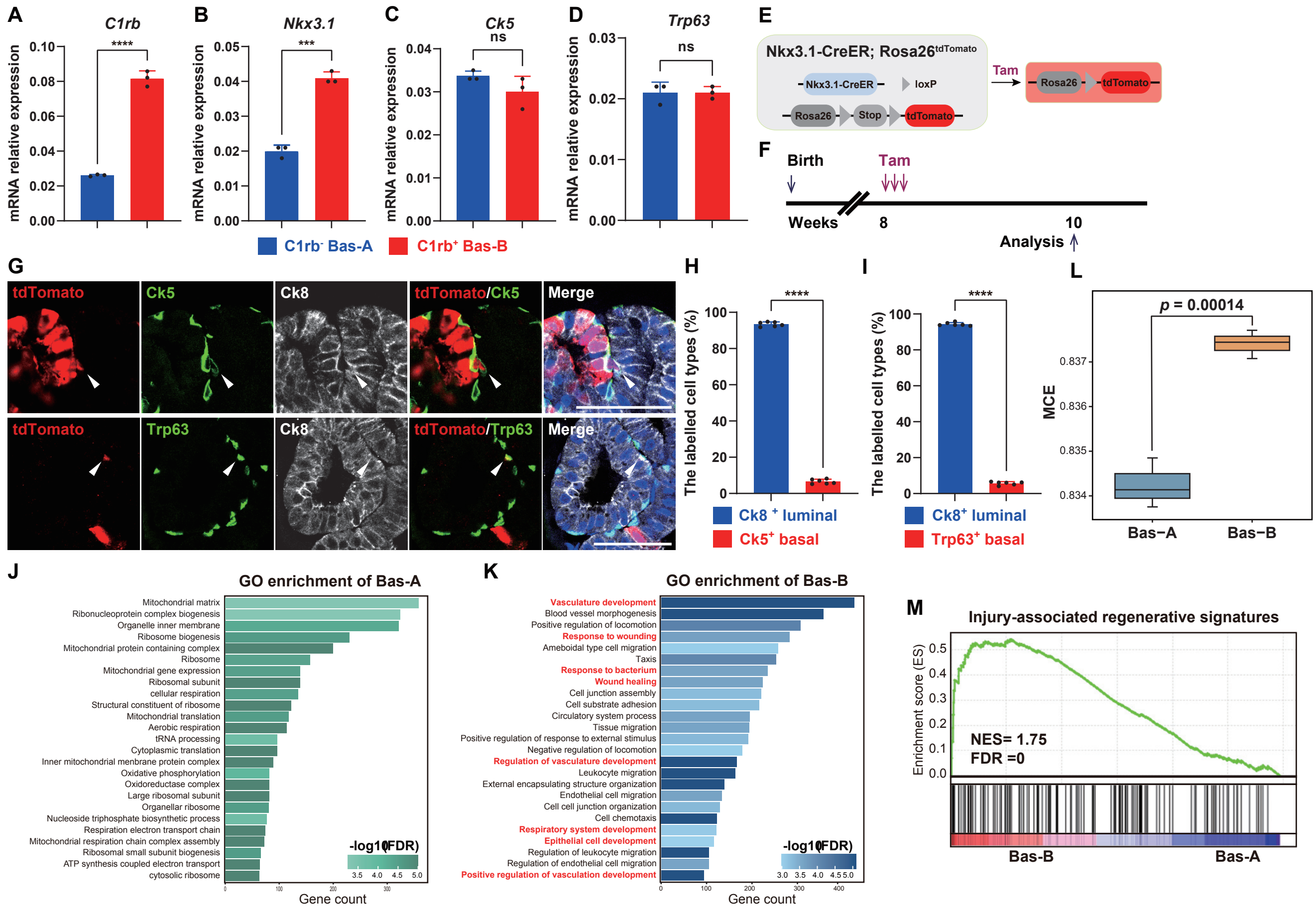

**Fig. S5. Isolation and characterization of mouse prostate Basal-A and Basal-B cells.** (**A** to **D**) Gene expression levels of *C1rb* (A), *Nkx3.1* (B), *Ck5* (C) and *Trp63* (D) in isolated C1rb$^-$ Basal-A and C1rb$^+$ Basal-B cells by qRT–PCR ($n = 3$ repeats per group). (**E**) Construction strategy of NT (*Nkx3.1-CreER; Rosa26-loxp-stop-loxp-tdTomato*) mice. (**F**) Schedule of labeling and tracing Nkx3.1-expressing cells in NT mice. (**G**) Immunofluorescence staining of tdTomato, Trp63, Ck5 and Ck8 in intact NT mouse prostates 2 weeks after Tam administration. (**H**) Percentage of Ck8$^+$ luminal cells and Ck5$^+$ basal cells among tdTomato$^+$ cells ($n = 6{,}754$) in NT mouse prostates ($n = 6$). (**I**) Percentage of Ck8$^+$ luminal cells and Trp63$^+$ basal cells among tdTomato$^+$ cells ($n = 7{,}214$) in NT mouse prostates ($n = 6$). (**J** and **K**) Top 25 significantly enriched (Methods; $p < 0.01$; $-\log_{10}$(P value)) gene ontology terms in the gene signature for the Basal-A (J) and Basal-B (K) subtype of bulk RNA analysis ($n = 3$). (**L**) Bar diagram shows MCE value of Basal-A and Basal-B cells. The P value is from a one-sided t test. (**M**) GSEA of injury-associated regenerative signatures in Basal-A and Basal-B cells. Scale bars, 50 µm. Data are shown as the mean ± SD. Data were analyzed by unpaired two-tailed Student's t test; ***$p < 0.001$, ****$p < 0.0001$ and ns, nonsignificant.

Fig. S6

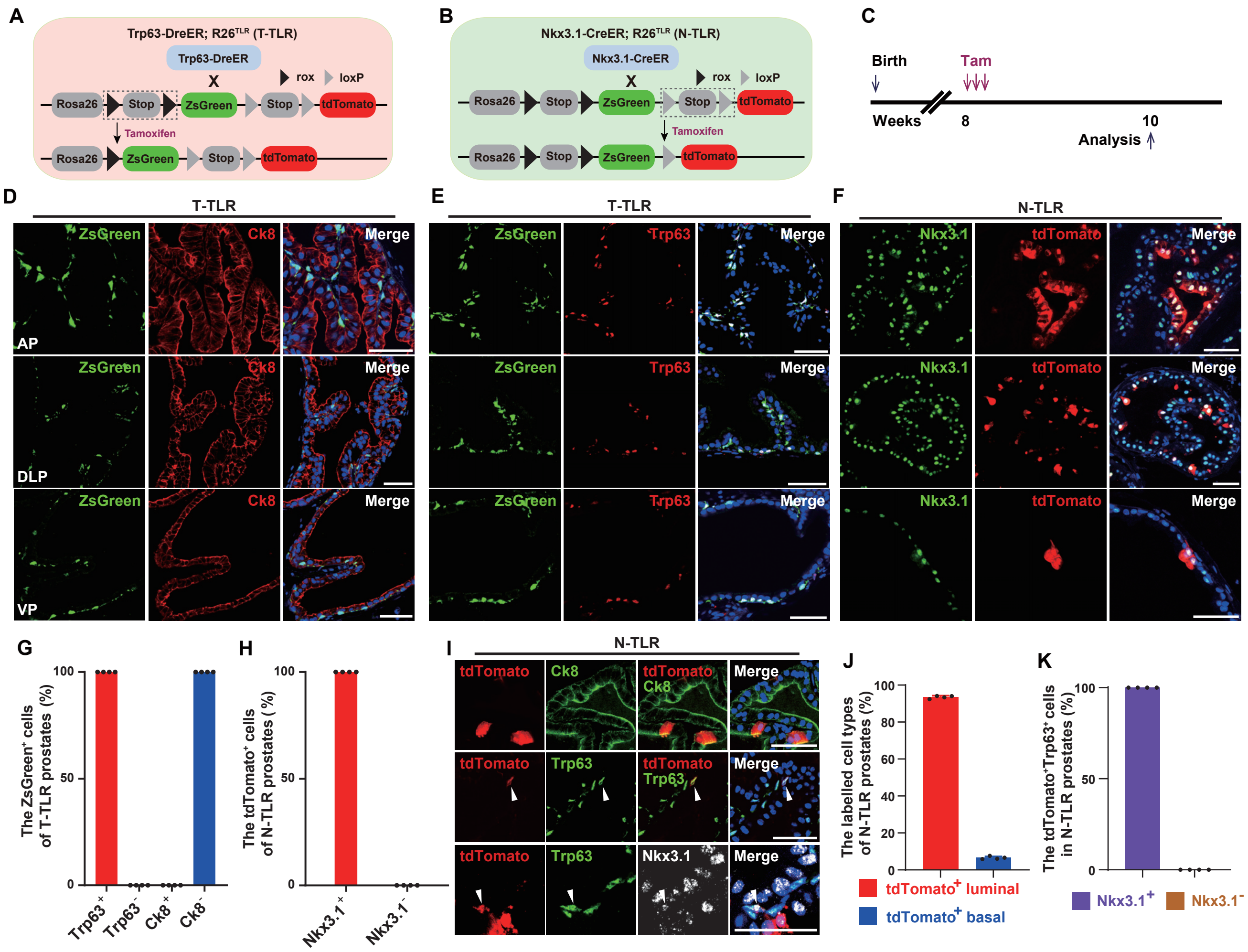

**Fig. S6. Trp63-expressing and Nkx3.1-expressing cell labeling.** (**A** and **B**) Construction strategy of T-TLR (*Trp63-DreER;R26-TLR*) mice (A) and N-TLR (*Nkx3.1-CreER;R26-TLR*) mice (B) for Trp63$^+$ and Nkx3.1$^+$ cell labeling, respectively. (**C**) Schedule of tamoxifen injection and analysis in T-TLR and N-TLR mouse prostates. (**D** to **F**) Immunofluorescence staining of Trp63, Nkx3.1, Ck8, ZsGreen and tdTomato in T-TLR (D and E) and N-TLR (F) mouse prostates. (**G**) Percentage of Trp63$^+$ ($n$ = 12,757), Trp63$^-$ ($n$ = 0), Ck8$^+$ ($n$ = 0), and Ck8$^-$ ($n$ = 15,300) cells among ZsGreen$^+$ cells in T-TLR mice ($n$ = 4). (**H**) Percentage of Nkx3.1$^+$ ($n$ = 14,438) and Nkx3.1$^-$ ($n$ = 0) cells among tdTomato$^+$ cells in N-TLR mice ($n$ = 4). (**I**) Immunofluorescence staining of Trp63, Nkx3.1, Ck8 and tdTomato in N-TLR mouse prostates. (**J**) Percentage of Ck8$^+$ luminal cells and Trp63$^+$ basal cells among tdTomato$^+$ cells ($n$ = 27,195) in N-TLR mice ($n$ = 4). (**K**) Percentage of Nkx3.1$^+$ cells ($n$ = 2,649) and Nkx3.1$^-$ cells ($n$ = 0) among tdTomato$^+$Trp63$^+$ cells in N-TLR mice ($n$ = 4). Scale bars, 50 µm. Data are shown as the mean ± SD.

Fig. S7

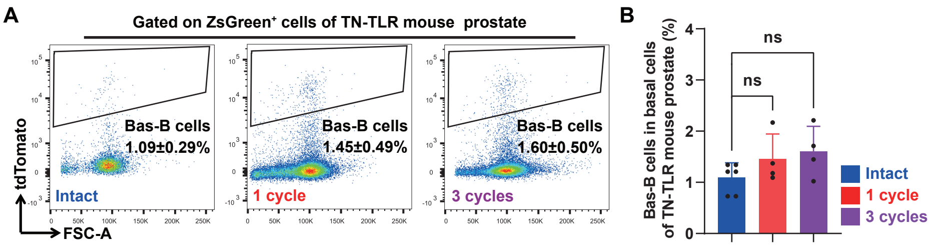

**Fig. S7. The percentage of Basal-B cell population during prostate regeneration.** (**A** and **B**) FACS plots (A) and a bar graph (B) showing the percentage of tdTomato$^{+}$ZsGreen$^{+}$ Basal-B cells among ZsGreen$^{+}$ basal cells of intact ($n$ = 6 mice) and regenerated TN-TLR mouse anterior prostates after 1 cycle ($n$ = 4 mice) or 3 cycles ($n$ = 4 mice) of prostate regression-regeneration.

Fig. S8

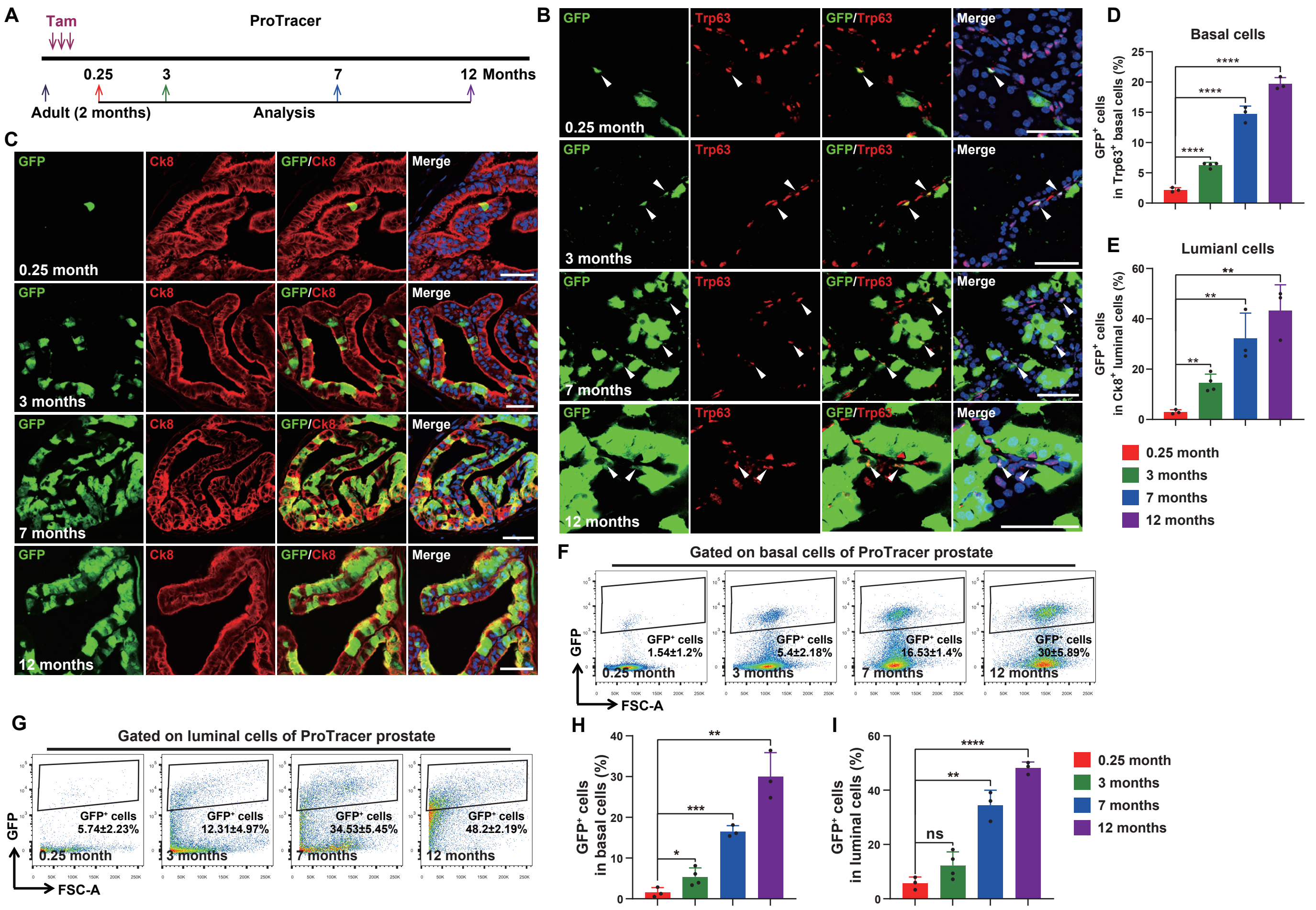

**Fig. S8. Prostate cell proliferation tracing during homeostasis using the ProTracer model.** (**A**) Schedule of tracing proliferated cells during prostate homeostasis in ProTracer mice. (**B** and **C**) Immunofluorescence staining of GFP and Trp63 for basal cells (B) and GFP and Ck8 for luminal cells (C) of ProTracer mouse prostates at 4 time points (1 week, 3 months, 7 months and 12 months) after tamoxifen injection. (**D** and **E**) Percentage of GFP$^+$ cells in Trp63$^+$ basal cells (D) or Ck8$^+$ luminal cells (E) of ProTracer mouse prostates at 4 time points (1 week ($n$ = 3 mice), 3 months ($n$ = 4 mice), 7 months ($n$ = 3 mice) and 12 months ($n$ = 3 mice)) after tamoxifen injection. (**F** to **I**) FACS plots and bar graphs showing the percentage of GFP$^+$ cells among Lin$^-$Epcam$^{high}$CD49f$^{high}$ basal cells (F and H) or Lin$^-$Epcam$^{high}$CD49f$^{low}$ luminal cells (G and I) of ProTracer mouse prostates at 4 time points (1 week ($n$ = 3 mice), 3 months ($n$ = 4 mice), 7 months ($n$ = 3 mice) and 12 months ($n$ = 3 mice)) after tamoxifen injection.

Fig. S9

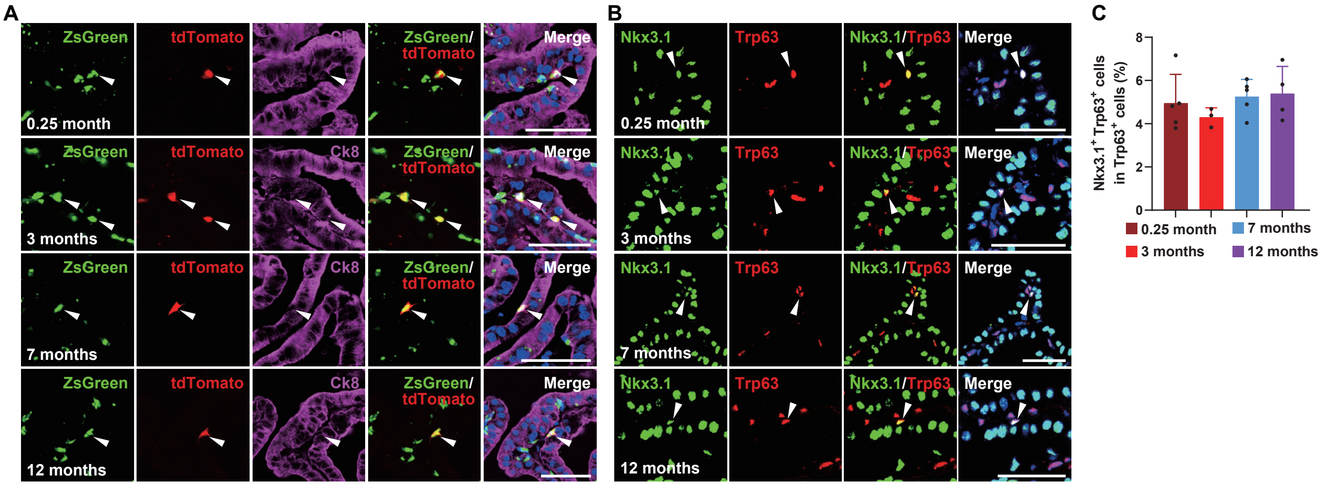

**Fig. S9. The percentage of Basal-B cells is stable during long-term prostate homeostasis.** (**A**) Immunofluorescence staining of ZsGreen and tdTomato in TN-TLR mouse prostates at 4 time points (1 week, 3 months, 7 months and 12 months) after tamoxifen injection. White arrows indicate ZsGreen$^+$tdTomato$^+$ Basal-B cells. (**B**) Immunofluorescence staining of Trp63 and Nkx3.1 in TN-TLR mouse prostates at 4 time points (1 week, 3 months, 7 months and 12 months) after vehicle injection. White arrows indicate Trp63$^+$Nkx3.1$^+$ Basal-B cells. (**C**) Percentage of Trp63$^+$Nkx3.1$^+$ Basal-B cells among Trp63$^+$ basal cells in TN-TLR mouse prostates at 4 time points (1 week (*n* = 5 mice), 3 months (*n* = 3 mice), 7 months (*n* = 5 mice) and 12 months (*n* = 4 mice)) after vehicle injection.

Fig. S10

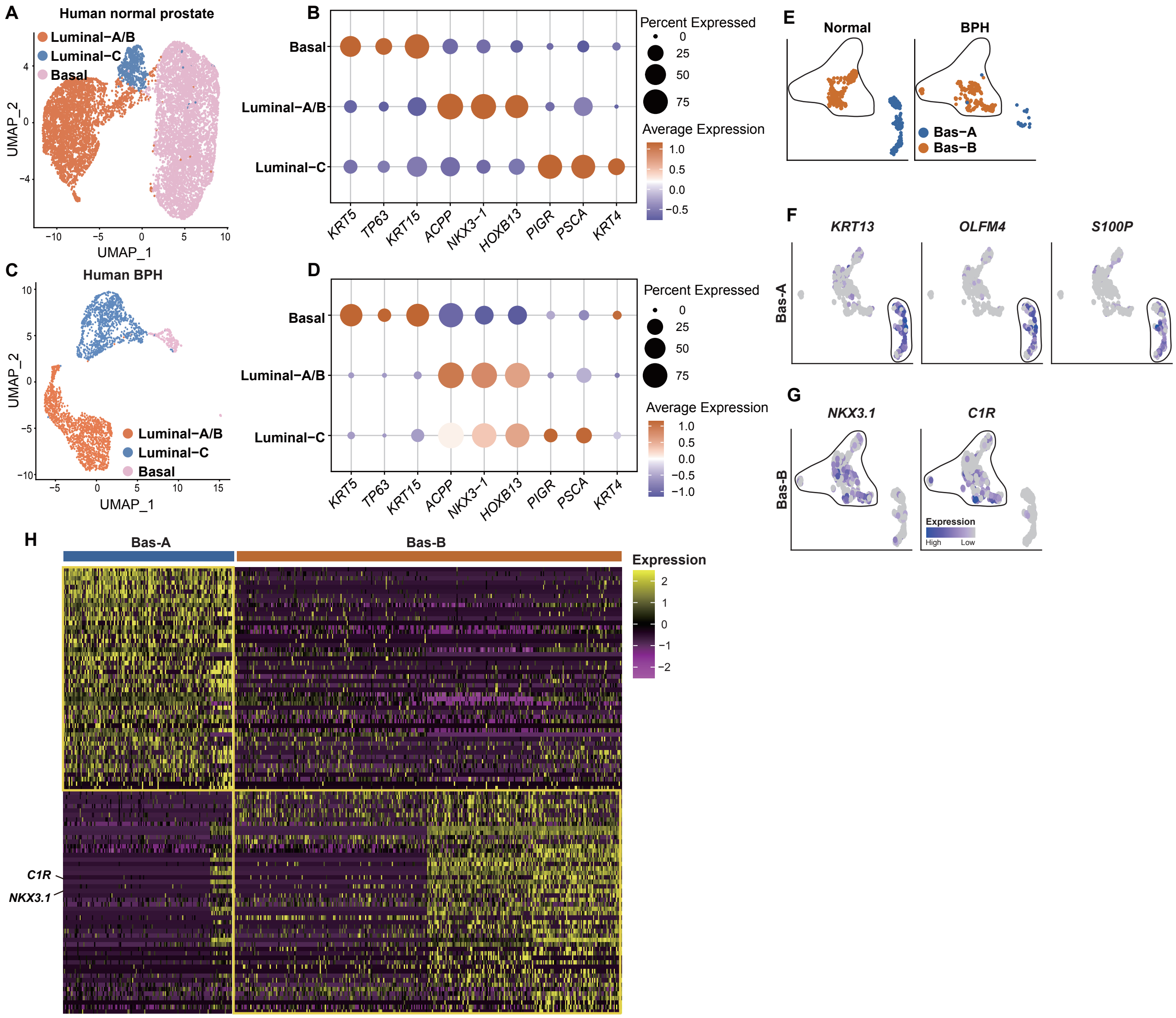

**Fig. S10. Integration analysis of basal cells from normal human prostate and BPH samples.** (**A**) Visualization of epithelial cell clustering results from normal human prostate samples. (**B**) The expression levels of representative signature genes across all epithelial subtypes in normal human prostate samples. (**C**) Visualization of epithelial cell clustering results from BPH samples. (**D**) The expression levels of representative signature genes across all epithelial subtypes in BPH samples. (**E**) Distribution of Basal-A and Basal-B in normal human prostate and BPH samples, respectively. (**F** and **G**) UMAP plots showing the expression levels of the basal signature genes *KRT5*, *TP63* and *KRT15* (F) and the Basal-B signature genes *NKX3.1* and *C1R* (G) across all basal cells. (**H**) Heatmap showing the expression levels of the top 50 signature genes across basal cells from normal human prostate and BPH samples.

Fig. S11

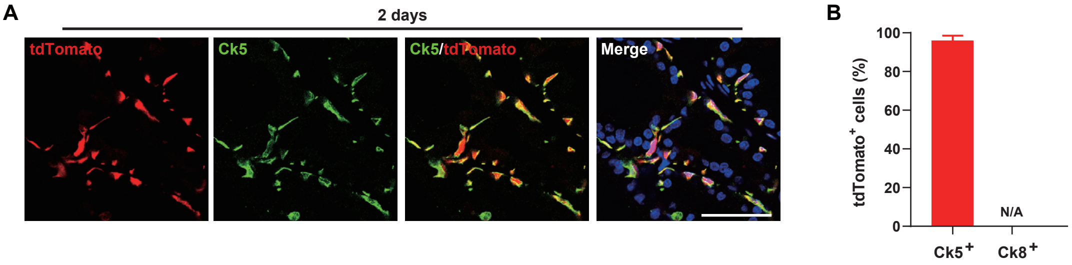

**Fig. S11. Basal cell labeling in K5CP mouse prostates.** (**A**) Immunofluorescence staining of Ck5 and tdTomato in K5CP mouse prostates 2 days after tamoxifen administration. (**B**) Percentage of tdTomato$^{+}$ cells among Ck5$^{+}$ basal cells ($n$ = 12,917) and Ck8$^{+}$ luminal cells ($n$ = 12,552) in K5CP mouse prostates ($n$ = 4). Scale bars, 50 µm. Data are shown as the mean ± SD.

Table S1. Primers. This table includes the primers used in this study (separate file).

The primers for genome typing used in this study:

| Primers | Sequences (5’ to 3’) | WT (bp) | Target gene (bp) |
|---|---|---|---|
| Nkx3.1-CreER forward 1 | AGCTATCCCTACTACCCCTACCT | 465 | 576 |
| Nkx3.1-CreER forward 2 | CTCACCGAAACCCAAGTCAAAA | | |
| Nkx3.1-CreER reverse 1 | CAGCCCCTTGTTGAATACG | | |
| Nkx3.1-CreER reverse 2 | GGAGCGGGCAGCAGTCACCTACA | | |
| Trp63-DreER forward 1 | ACAGCCACAGTACACGAACC | 518 | 806 |
| Trp63-DreER forward 2 | CTATTGCTTCCCGTATGGCTTTCA | | |
| Trp63-DreER reverse 1 | GCATATACAGCACCTCCTAA | | |
| Trp63-DreER reverse 2 | GTGTTGTAGGGGCTGGTGGACGAG | | |
| Rosa26-TLR forward 1 | TTGGAGGCAGGAAGCACTTG | 364 | 251 |
| Rosa26-TLR forward 2 | ACCATCGTGGAACAGTACGAGC | | |
| Rosa26-TLR reverse 1 | CCGACAAAACCGAAAATCTGTG | | |
| Rosa26-TLR reverse 2 | TTGGTCACCTTCAGCTTGGC | | |
| Ck5-CreER forward 1 | GCCTTGGCCCACGCTTCA | 458 | 610 |
| Ck5-CreER forward 2 | GGCCCACGCTTCACCAG | | |
| Ck5-CreER reverse 1 | GTGGCTTACATTCTGCAACATTTT | | |
| Ck5-CreER reverse 2 | GGATCCGCCGCATAACCAGT | | |
| Rosa26-LSL-tdTomato forward 1 | AAG GGA GCT GCA GTG GAG TA | 297 | 196 |
| Rosa26-LSL-tdTomato forward 2 | CCG AAA ATC TGT GGG AAG TC | | |
| Rosa26-LSL-tdTomato reverse 1 | GGC ATT AAA GCA GCG TAT CC | | |
| Rosa26-LSL-tdTomato reverse 2 | CTG TTC CTG TAC GGC ATG G | | |
| Ck8-CreER forward 1 | GCGGTCTGGCAGTAAAAACTATC | 324 | 100 |
| Ck8-CreER forward 2 | CTAGGCCACAGAATTGAAAGATCT | | |
| Ck8-CreER reverse 1 | GTGAAACAGCATTGCTGTCACTT | | |
| Ck8-CreER reverse 2 | GTAGGTGGAAATTCTAGCATCATCC | | |
| Pten$^{flox/flox}$ forward | TGTTTTTGACCAATTAAAGTAGGCTGTG | 350 | 490 |
| Pten$^{flox/flox}$ reverse | AAAAGTTCCCCTGCTGATGATTTGT | | |
| Rosa26-DreER forward 1 | CGTGCTGGTTATTGTGCTGTCTC | 364 | 143 |
| Rosa26-DreER forward 2 | TTGGAGGCAGGAAGCACTTG | | |
| Rosa26-DreER reverse 1 | TACTCCTTGCCGATGTTCCTCAGG | | |
| Rosa26-DreER reverse 2 | CCGACAAAACCGAAAATCTGTG | | |
| Rosa26-LSL-GFP forward 1 | CAGCGACTTCTTCATCCAGAGC | 297 | 622 |
| Rosa26-LSL-GFP forward 2 | AAGGGAGCTGCAGTGGAGTA | | |
| Rosa26-LSL-GFP reverse 1 | AAAGCAGCGTATCCACATAGCG | | |
| Rosa26-LSL-GFP reverse 2 | CCGAAAATCTGTGGGAAGTC | | |
| Ki67-CrexER forward 1 | GGGCTCTACTTCATCGCATTCC | 475 | 312 |
| Ki67-CrexER reverse 1 | ATCTGGTTCCCTGGATGGTTG | | |
| Ki67-CrexER reverse 2 | TTGGCGTCTGAAGAGAGTATGACC | | |

| | | | |
|---|---|---|---|
| Nkx3.1-CrexER forward 1 | GAAAGCAGCTGTCGGAAGAC | 465 | 739 |
| Nkx3.1-CrexER forward 2 | CTCACCGAAACCCAAGTCAAAA | | |
| Nkx3.1-CrexER reverse 1 | TAAGCAATCCCCAGAAATGC | | |
| Nkx3.1-CrexER reverse 2 | GGAGCGGGCAGCAGTCACCTACA | | |
| The qRT–PCR primers used in this study: | | | |
| Primers | Sequence (5' - 3') | | |
| Actb-qPCR-F | CATTGCTGACAGGATGCAGAAGG | | |
| Actb-qPCR-R | TGCTGGAAGGTGGACAGTGAGG | | |
| Krt5-qPCR-F | GAACAGAGGCTGAGTCCTGGTA | | |
| Krt5-qPCR-R | TCTCAGCCTCTGGATCATTCGG | | |
| Trp63-qPCR-F | GTATCGGACAGCGCAAAGAACG | | |
| Trp63-qPCR-R | CTGGTAGGTACAGCAGCTCATC | | |
| C1rb-qPCR-F | ATCAGGCGCTACTGTCCTTCAC | | |
| C1rb-qPCR-R | TGTGGTGATGTAGCGGAAGTCC | | |
| Nkx3.1-qPCR-F | CGCGGAGACACCGACTGAACC | | |
| Nkx3.1-qPCR-R | TTCTGTGGCTGCTTGGTGACCT | | |

Table S2. Antibodies. This table includes the antibodies used in this study (separate file).

| The primary antibodies for immunofluorescence used in this study: | | | |
|---|---|---|---|
| Antigen | Dilution | Supplier | Catalog# |
| Nkx3.1 (for mouse prostates) | 1:1000 | AthenaES | 315 |
| NKX3.1 (for human prostates) | 1:250 | Cell Signaling Technology | 83700S |
| Ck8 | 1:1000 | Covance | MMS-162-P |
| Ck8 | 1:100 | Developmental Studies Hybridoma Bank | TROMA-I |
| Ck5 | 1:1000 | Covance | PRB-160P |
| Trp63 | 1:100 | Abcam | ab735 |
| Trp63 | 1:250 | Abcam | ab124762 |
| CD45 | 1:500 | Abcam | ab10558 |
| ZsGreen | 1:500 | Clontech | 632598 |
| DsRed | 1:250 | Takara | 632496 |
| The antibodies for FACS used in this study: | | | |
| Antibody | Dilution | Supplier | Catalog# |
| CD45-APC | 1:500 | Invitrogen | 17-0451-83 |
| Ter119-APC | 1:500 | Invitrogen | 17-5921-82 |
| CD31-APC | 1:500 | Invitrogen | 17-0311-82 |
| Epcam-APC/Cy7 | 1:500 | Biolegend | 118218 |
| CD49f-AF700 | 1:500 | Invitrogen | 56-0495-82 |
| C1rb | 1:100 | Invitrogen | PA5-109463 |

The secondary antibodies used in this study:

| Antibody | Dilution | Supplier | Catalog# |
|---|---|---|---|
| Goat anti-Mouse IgG, Alexa Fluor 647 | 1:500 | Invitrogen | A21236 |
| Goat anti-rat IgG, Alexa Fluor 647 | 1:500 | Invitrogen | A21247 |
| Goat anti-rabbit IgG, Alexa Fluor 647 | 1:500 | Invitrogen | A21244 |
| Goat anti-Mouse IgG, Alexa Fluor 488 | 1:500 | Invitrogen | A11029 |
| Goat anti-rabbit IgG, Alexa Fluor 488 | 1:500 | Invitrogen | A11034 |
| Goat anti-Mouse IgG, Alexa Fluor 594 | 1:500 | Invitrogen | A11005 |
| Goat anti-rabbit IgG, Alexa Fluor 594 | 1:500 | Invitrogen | A11037 |

**Table S3**. Signature genes of basal cell subtypes in mouse prostates (separate file).

**Table S4**. Signature genes of basal cell subtypes in human prostates (separate file).

**Table S5**. Signature genes of Basal-A cells and Basal-B cells from bulk RNA-seq data (separate file).

**Table S6**. Enrichment results of Basal-A cells and Basal-B cells based on GSEA (separate file).
